# Effector-mediated subversion of proteasome activator (PA)28αβ enhances host defense against *Legionella pneumophila* under inflammatory and oxidative stress conditions

**DOI:** 10.1101/2021.12.27.473672

**Authors:** Tshegofatso Ngwaga, Deepika Chauhan, Abigail G. Salberg, Stephanie R. Shames

## Abstract

*Legionella pneumophila* is a natural pathogen of amoebae that causes Legionnaires’ Disease in immunocompromised individuals via replication within macrophages. *L. pneumophila* virulence and intracellular replication hinges on hundreds of Dot/Icm-translocated effector proteins, which are essential for biogenesis of the replication permissive *Legionella*-containing vacuole (LCV). However, effector activity can also enhance mammalian host defense via effector-triggered immunity. The *L. pneumophila* effector LegC4 is important for virulence in amoebae but enhances host defense against *L. pneumophila* in the mouse lung and, uniquely, within macrophages activated with either tumor necrosis factor (TNF) or interferon (IFN)-γ. The mechanism by which LegC4 potentiates cytokine-mediated host defense in macrophages is unknown. Here, we found that LegC4 enhances cytokine-mediated phagolysosomal fusion with *Legionella*-containing vacuole (LCV) and binds host proteasome activator (PA)28α, which forms a heterooligomer with PA28β to facilitate ubiquitin-independent proteasomal degradation of oxidant-damaged (carbonylated) proteins. We found that oxidative stress was sustained in the presence of LegC4 and that the LegC4 restriction phenotype was relieved in PA28αβ-deficient macrophages and in the lungs of mice *in vivo*. Our data also show that oxidative stress is sufficient for LegC4-mediated restriction in macrophages producing PA28αβ. PA28αβ has been traditionally associated with antigen presentation; however, our data support a novel mechanism whereby effector-mediated subversion of PA28αβ enhances cell-autonomous host defense against *L. pneumophila* under inflammatory and oxidative stress conditions. This work provides a solid foundation to evaluate induced proteasome regulators as mediators of innate immunity.

**Author Summary:** Pro-inflammatory cytokines induce antimicrobial host defense pathways within macrophages to control intracellular pathogens. We discovered that the *Legionella pneumophila* effector protein LegC4 potentiates pathogen clearance within cytokine-activated macrophages. Here, we show a central role for host proteasome activator (PA)28αβ in LegC4 restriction. PA28αβ is upregulated by cytokine signaling and under oxidative stress conditions to facilitate proteasomal degradation of oxidant-damaged proteins. We found that LegC4 binds PA28αβ and that LegC4 restriction was lost in PA28αβ-deficient (*Psme1/2*^-/-^) macrophages and a mouse model of Legionnaires’ Disease *in vivo*. Furthermore, oxidative stress was sustained in the presence of LegC4 and was sufficient for LegC4 restriction in PA28αβ-producing cells macrophages. Finally, we found that *L. pneumophila* replication was attenuated within PA28αβ-deficient macrophages irrespective of LegC4. These findings support a model whereby subversion of host proteostasis machinery triggers cell-autonomous host defense within macrophages under inflammatory and oxidative stress conditions.

## Introduction

Many intracellular bacterial pathogens cause disease by replicating within host macrophages [1]. However, macrophages activated by T_H_1 proinflammatory cytokines, such as tumor necrosis factor (TNF) and interferon (IFN)-γ, are potently and broadly microbicidal. However, activated macrophages and inflamed tissues are in a state of altered protein homeostasis (proteostasis), owing in part to robust production of reactive oxygen species (ROS), which cause collateral damage to cellular proteins. To cope with this oxidative stress, cytokine signaling and ROS upregulate proteasome activator (PA)28αβ, which engages proteolytic 20S proteasome core particles (CPs) to facilitate rapid ubiquitin-independent degradation of oxidant-damaged proteins [2]. Impaired proteasome activity and proteotoxic stress trigger compensatory upregulation of lysosomal degradation pathways, which are central to cell-autonomous immunity within activated macrophages [3]. PA28αβ has been traditionally associated with antigen presentation and adaptive immunity [4,5]; however, a role for PA28αβ in cell-autonomous antimicrobial innate host defense has not been established.

*Legionella pneumophila* is a natural pathogen of amoebae and accidental human pathogen that causes Legionnaires’ Disease pneumonia in immunocompromised individuals that results from bacterial replication within alveolar macrophages [6]. *L. pneumophila* replication within macrophages is a direct consequence of extensive co-evolution with natural host amoebae [7–9]; however, *L. pneumophila* rarely transmits person-to-person and is rapidly cleared by acute pathogen-centric innate immune responses in healthy hosts [10–12]. To establish its intracellular niche, the *Legionella*-containing vacuole (LCV), *L. pneumophila* translocates hundreds of virulence factors, called effector proteins, into infected host cells by a Dot/Icm type IV secretion system (T4SS) [6,13,14]. Effectors are essential for virulence, but detection of their activity within host cells can induce pro-inflammatory cytokine production via effector-triggered immunity (ETI) [10,15–17].

Pro-inflammatory cytokines activate and potentiate the antimicrobial capacity of infected macrophages and are central to host defense against *L. pneumophila* [18,19]. Thus, *L. pneumophila* is a well-established model pathogen to study mechanisms of pathogen-centric and effector-mediated antimicrobial innate immunity.

We discovered that the *L. pneumophila* effector LegC4 induces a unique mechanism of effector-mediated host defense by potentiating the antimicrobial activity of activated macrophages [20]. LegC4 host defense by confers a fitness disadvantage on *L. pneumophila* in a mouse model of Legionnaires’ Disease, likely resulting from elevated levels of pro-inflammatory cytokines in the lung since a growth defect is only observed within cultured macrophages when activated with exogenous TNF or IFN-γ [20,21]. The evolutionary basis for retention of LegC4 by *L. pneumophila* stems from its importance for replication within its natural host, *Acanthamoeba castellanii* [21]. Thus, LegC4 confers quantifiable opposing phenotypes in natural (protozoa) and ‘accidental’ (mammalian) hosts, which is noteworthy since the majority of *L. pneumophila* effectors are functionally redundant in laboratory models and have no obvious growth phenotypes [21,22]. The specific augmentation of cytokine-mediated restriction by LegC4 represents a potentially novel mechanism of ETI, but how LegC4 functions and potentiates host defense against *L. pneumophila* is unknown.

Here, we investigated the mechanism of LegC4 restriction within macrophages and found a central role for PA28αβ and oxidative stress. We found a central role for PA28αβ, phagolysosomal fusion, and oxidative stress in LegC4-mediated restriction. LegC4 enhanced cytokine-dependent phagolysosomal fusion with LCVs and perpetuated oxidative stress. LegC4 bound host PA28α *in vitro* and changed the subcellular localization of PA28α within infected and transfected cells. LegC4 restriction phenotypes were relieved in PA28αβ-deficient (*Psme1/2*^-/-^) BMDMs and the fitness advantage associated with Δ*legC4* mutant bacteria *in vivo* was abolished in *Psme1/2*^-/-^ mice. The abundance of oxidant-damaged proteins was increased in the presence of LegC4 and oxidative stress was sufficient to impair replication of LegC4-producing of *L. pneumophila* in BMDMs producing PA28αβ. Importantly, *L. pneumophila* replication was attenuated within *Psme1/2*^-/-^ BMDMs irrespective LegC4, suggesting that impaired PA28αβ activity is sufficient to augment host defense in macrophages and is the mechanism driving LegC4 restriction. Overall, our data support a model whereby PA28α targeting by a translocated effector protein perturbs proteostasis and enhances cell-autonomous restriction of *L. pneumophila* within activated macrophages. These data are the first to uncover a role for PA28αβ in innate host defense against an intracellular bacterial pathogen and reveal a novel mechanism by which the antimicrobial activity of macrophages can be augmented under inflammatory and oxidative stress conditions.

## Results

### LegC4 confers a fitness disadvantage on *L. pneumophila* in cytokine-activated macrophages and in a mouse model of Legionnaires’ Disease

We previously reported that LegC4 confers a fitness disadvantage on *L. pneumophila* in intranasal C57Bl/6 mouse models of Legionnaires’ Disease (LD) and within cytokine-activated macrophages [20,21]. C57Bl/6 mice are restrictive to *L. pneumophila* due to flagellin (FlaA)-mediated activation of the NAIP5/NLRC4 inflammasome [23,24]. Thus, to study FlaA-independent host responses within C57Bl/6 mice, we leverage FlaA-deficient *L. pneumophila* (Δ*flaA*) strains. We found that *L. pneumophila* Δ*flaA*Δ*legC4* (herein called ΔΔ) mutant strains outcompeted parental Δ*flaA* strains in the lungs of C57Bl/6 wild-type (WT) mice [20] but a growth phenotype for *L. pneumophila* ΔΔ was only observed within cultured mouse primary bone marrow-derived macrophages (BMDMs) activated with exogenous recombinant (r)TNF or rIFN-γ [20]. However, autocrine and paracrine TNF produced by cultured BMDMs is sufficient to attenuate *L. pneumophila* strains induced to overexpress *legC4* from a complementing multicopy plasmid in naïve WT BMDMs [20]. Importantly, this phenotype was specific to intracellularly growing bacteria since *L. pneumophila* growth in broth *in vitro* is unaffected by overexpression of *legC4* [20]. Thus, LegC4 confers a growth defect on *L. pneumophila* in a dose-dependent manner within cytokine-activated macrophages.

We confirmed our previous results and found that overexpression of *legC4* was sufficient to impair *L. pneumophila* replication within WT BMDMs since *L. pneumophila* ΔΔ (p*legC4*) replication was significantly attenuated compared to both the *L. pneumophila* Δ*flaA* parental strain and *L. pneumophila* ΔΔ harboring an empty plasmid vector (pEV) (**Fig 1A**). Impaired replication was dependent on TNF signaling since overexpression of *legC4* did not impair growth within BMDMs unable to produce TNF (*Myd88*^-/-^; **Fig 1B**) or signal through TNFR1 (*Tnfr1*^-/-^; **Fig 1C**). IFN-γ signaling is sufficient for LegC4 restriction [20]; however, the LegC4 restriction phenotype was retained within *Ifngr1*^-/-^ BMDMs (**Fig 1D**), indicating that the relatively low levels of IFN-γ secreted by macrophages are not sufficient for restriction. Importantly, we found that LegC4 restriction is not specific to mouse macrophages since both *L. pneumophila* Δ*flaA* and ΔΔ (p*legC4*) were impaired for replication compared to the ΔΔ (pEV) strain in PMA-differentiated human THP-1 macrophages (**Fig 1E**). Thus, LegC4 confers a dose-dependent virulence defect on *L. pneumophila* within mouse and human macrophages.

**Figure 1.**
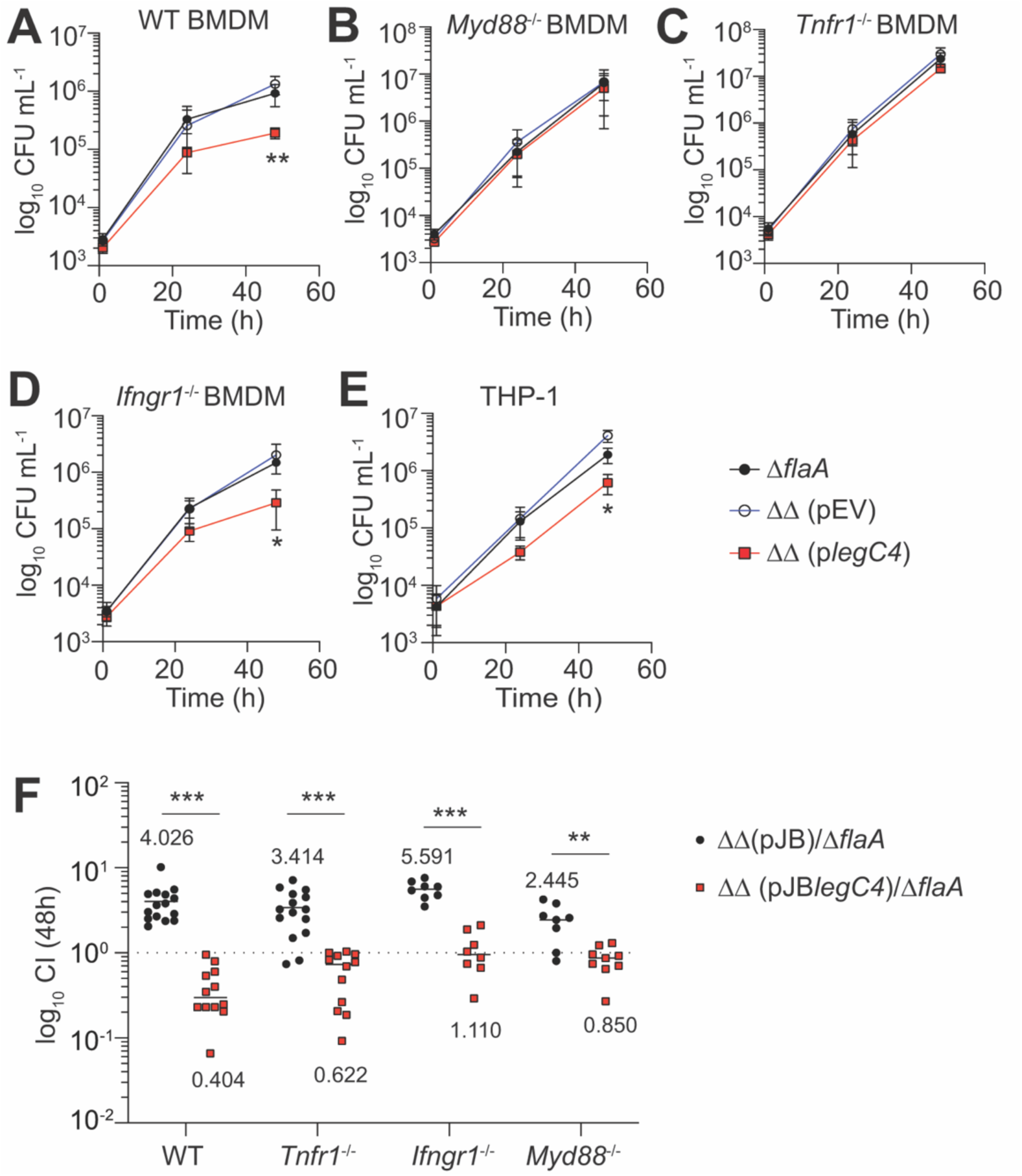
LegC4 attenuates *L. pneumophila* replication within cytokine-activated macrophages and the mouse lung. Growth of indicated *L. pneumophila* strains within **(A)** WT BMDMs, **(B)** *Myd88*^-/-^ BMDMs, **(C)** *Tnfr1*^-/-^ BMDMs, or **(D)** *Ifngr1*^-/-^ BMDMs, or **(E)** THP-1 cells over 48 h. Plasmid expression of *legC4* was induced with 1 mM IPTG. Data are shown as mean ± s.d. of pooled results of at least two independent experiments with triplicate wells per condition in each experiment. Asterisks denote statistical significance by Students’ *t*-test (\**P*<0.05, \*\**P*<0.01). ΔΔ: Δ*flaA*Δ*legC4*. **(F)** Competitive index (CI) values from the lungs of mice infected with a 1:1 mixture of *L. pneumophila* ΔΔ (pJB*legC4*) or Δ*flaA*Δ*legC4* (pJB) with *L. pneumophila* Δ*flaA* at 48 h post-infection. Each symbol represents an individual animal and data are pooled from at least two independent experiments and mean CI values are shown. Asterisks denote statistical significance by two-Way ANOVA (\*\*\**P*<0.0001, \*\**P*<0.01). ΔΔ: Δ*flaA*Δ*legC4*

Cytokine signaling drives LegC4 growth defects in cultured BMDMs, but the role of cytokine signaling *in vivo* is unclear. We evaluated bacterial fitness by calculating competitive index values for mice infected with a 1:1 mixture of *L. pneumophila* Δ*flaA* and either *L. pneumophila* ΔΔ (pEV) or ΔΔ (p*legC4*) for 48 h (see *Materials & Methods*). We confirmed our previous results showing that LegC4 confers a dose-dependent fitness defect on *L. pneumophila* in the lungs of WT mice since *L. pneumophila* Δ*flaA* was outcompeted by *L. pneumophila* ΔΔ (pEV) but was able to outcompete *L. pneumophila* ΔΔ (pJB*legC4*; **Fig 1F**) [20]. We also confirmed that loss of TNFR1 is insufficient to abolish the *legC4* mutant fitness advantage (**Fig 1F**) and obtained similar results in the lungs of IFNGR1- and MyD88-deficient mice (**Fig 1F**). However, the fitness phenotype associated with loss LegC4 was less pronounced in the lungs of *Myd88*^-/-^ mice compared to WT, *Tnfr1*^-/-^ and *Ifngr1*^-/-^ mice (**Fig 1F**), which is likely due to the relatively low levels of MyD88-indpendent IFN-γ produced in the *L. pneumophila*-infected lung [25]. Furthermore, the fitness disadvantage associated with overexpression of *legC4* was less pronounced in the lungs of MyD88-, IFNGR1- and TNFR1-deficient mice compared to WT mice (**Fig 1F**). These data support our previous work showing that LegC4 augments redundant cytokine-mediated pathogen restriction mechanisms [20]. Overexpression of *legC4* is clearly not necessary to observe a phenotype; however, based on the narrow phenotypic window in BMDMs activated with exogenous cytokine, this infection model is useful for our goal to define how LegC4 potentiates cytokine-mediated restriction within macrophages

### LegC4 induces robust phagolysosomal fusion with LCVs BMDMs

LegC4 attenuates *L. pneumophila* replication within cytokine-activated BMDMs; however, the mechanism of bacterial restriction is unknown. Cytokine signaling enhances phagolysosomal fusion with intracellular pathogens [26–29]; thus, we tested the hypothesis that LegC4 restriction involves lysosomal targeting of *L. pneumophila*. Virulent *L. pneumophila* establish a replication-permissive compartment called the *Legionella*-containing vacuole (LCV) by blocking endocytic maturation of their phagosome [30,31]. Avirulent Dot/Icm-deficient *L. pneumophila* (Δ*dotA*) are unable to establish a replicative LCV and phagosomes rapidly mature to endolysosomes, as evidenced by presence of lysosomal membrane protein LAMP1 [30]. We initially tested the hypothesis that LegC4 induces phagolysosomal fusion with LCVs by blinded immunofluorescence scoring of LAMP1^+^ LCVs within BMDMs infected with *L. pneumophila* strains for 9h. Using our *legC4* overexpression model, we observed robust LegC4-dependent phagolysosomal fusion with significantly more LAMP1^+^ *L. pneumophila* ΔΔ (p*legC4*) LCVs were noted relative to both *L. pneumophila* Δ*flaA* or ΔΔ (pEV) control strains (**Fig 2A**). Strikingly, the abundance of LAMP1^+^ LCVs within WT BMDMs infected with *L. pneumophila* ΔΔ (p*legC4*) did not differ from cells infected with the avirulent Δ*dotA* control strain (**Fig 2A**). Moreover, robust LegC4-mediated phagolysosomal fusion with LCVs was significantly suppressed within TNFR1-deficient BMDMs (**Fig 2A**) since *Tnfr1*^-/-^ BMDMs infected with *L. pneumophila* ΔΔ (p*legC4*) harbored significantly fewer LAMP1^+^ LCVs than those infected with the Δ*dotA* control strain (**Fig 2A**). Interestingly, loss of TNFR1-mediated signaling did not fully abolish phagolysosomal fusion with *L. pneumophila* ΔΔ (p*legC4*) LCVs, but our growth curve data suggest that this is not sufficient for detectable differences in intracellular replication (**Fig 1C**).

**Figure 2.**
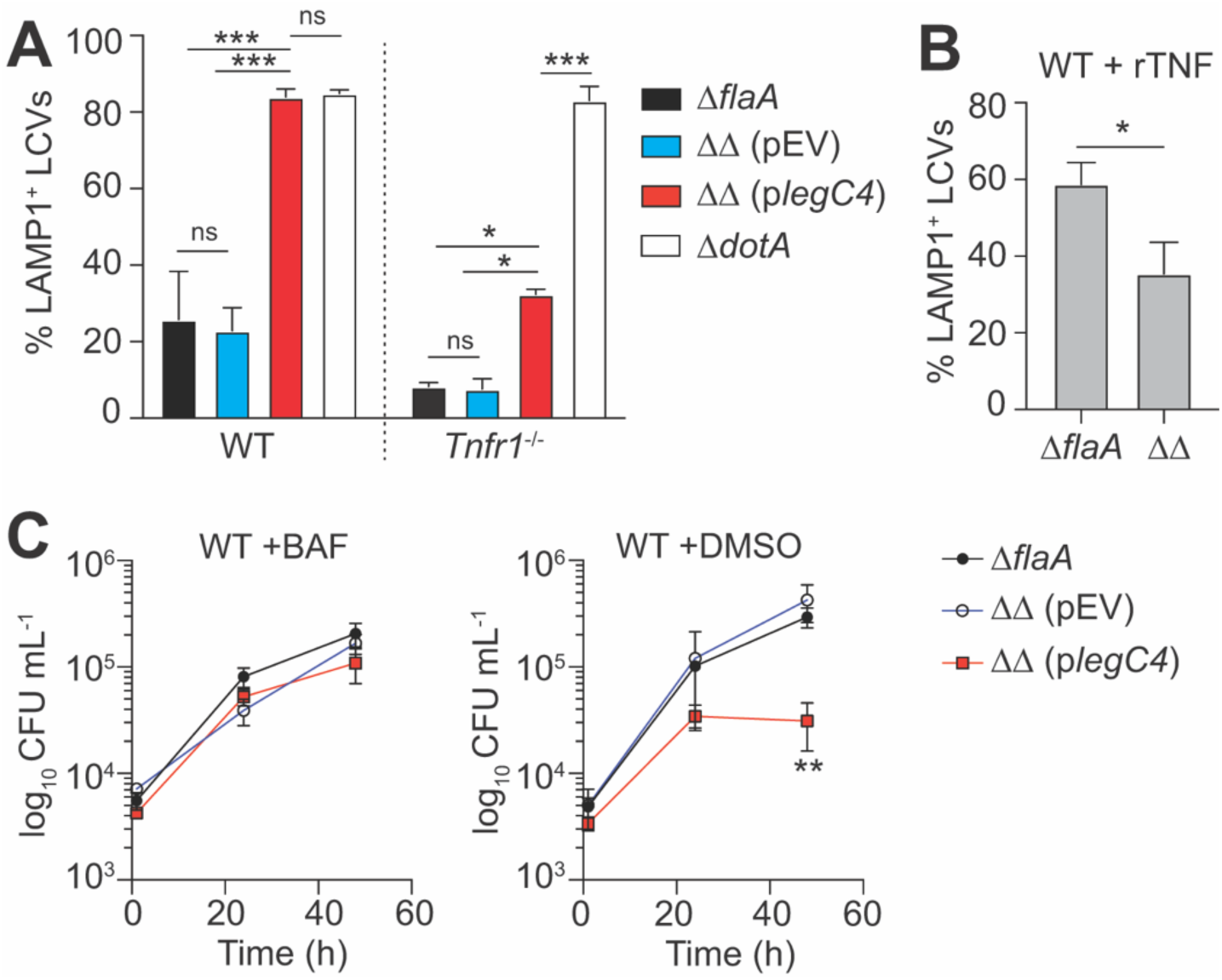
LegC4 enhances phagolysosomal fusion with LCVs and restriction is suppressed by inhibition of the vacuolar (v)ATPase. Abundance of LAMP1+ LCVs within **(A)** naïve WT or *Tnfr1*^-/-^ BMDMs infected with *L. pneumophila* strains for 9h; or **(B)** rTNF-treated WT BMDMs (25 ng mL^-1^) infected with *L. pneumophila* strains for 6 h. Infections were performed in triplicates and LAMP1 abundance was scored blind (*n*=100 cells per coverslip) by immunofluorescence microscopy (*n*=300 per condition per experiment). Plasmid expression of *legC4* was induced with 1 mM IPTG. Data shown are mean ± s.e.m. of pooled results from two independent experiments (*n*=600 total per condition). Asterisks denote statistical significance by two-way ANOVA (\**P*<0.05; \*\**P*<0.01; ns, not significant). **(C)** Growth of *L. pneumophila* strains within WT BMDMs treated with 10 nM bafilomycin A1 (BAF) (left panel) or DMSO (vehicle control; right panel). Plasmid expression of *legC4* was induced with 1 mM IPTG. Data shown are mean ± s.d. of pooled results from three independent experiments with triplicate samples per experiment. Asterisks denote statistical significance by Students’ *t*-test (\*\**P*<0.01). ΔΔ: Δ*flaA*Δ*legC4*.

Since endogenous LegC4 suppresses *L. pneumophila* replication within BMDMs activated with exogenous rTNF [20], we hypothesized that endogenous LegC4 would also be sufficient to induce phagolysosomal fusion with LCVs in rTNF-treated BMDMs. Indeed, more LAMP1^+^ LCVs were present in rTNF-treated BMDMs infected with *L. pneumophila* Δ*flaA* compared to *L. pneumophila* ΔΔ (**Fig 2B**). Together, these data show that LegC4 augments phagolysosomal fusion with LCVs in cytokine-activated BMDMs.

### Pharmacological inhibition of the H^+^-type vacuolar (v)ATPase abolishes LegC4 restriction in BMDMs

We subsequently tested whether lysosomal acidification contributes to LegC4-mediated restriction by quantifying *L. pneumophila* replication within BMDMs treated with Bafilomycin A1 (BAF). BAF is a pharmacological inhibitor of the H^+^-type vacuolar (v)-ATPase, which drives vacuolar acidification and phagolysosomal fusion [32]. Thus, we tested a role for vATPase activity in LegC4 restriction by quantifying *L. pneumophila* replication within BAF-treated BMDMs. LCV acidification is important for late stages of the *L. pneumophila* lifecycle [33]; however, intracellular replication is minimally impacted within BMDMs treated with a low concentration (≤12.5 nM) of BAF [34]. We found that *L. pneumophila* ΔΔ (p*legC4*) replication did not differ from *L. pneumophila* Δ*flaA* and ΔΔ (pEV) strains within BMDMs treated with 10 nM BAF (**Fig 2C, left**). However, as expected, *L. pneumophila* ΔΔ (p*legC4*) replication was significantly attenuated compared to Δ*flaA* and ΔΔ (pEV) within vehicle (DMSO)-treated BMDMs (**Fig 2C, right**). These data further support a role for lysosomes in LegC4-mediated restriction.

### LegC4 interacts with host proteasome activator (PA)28α

To gain insight into the mechanism of LegC4 function, we identified LegC4 binding partners by yeast two-hybrid (Y2H) using LegC4 as bait and 30,000 clones of fragmented genomic (g)DNA from mouse splenocytes as prey (Hybrigenics, Inc., see *Materials & Methods*). Our screen revealed interactions between LegC4 and several regulators of eukaryotic proteasomes, including proteasome activator (PA)28α, PA28γ, and Ecm29 (**Table S1**). The highest-confidence interactor was PA28α with 28 individual clones, all of which encoded a minimal overlapping region in the PA28α C-terminal domain (amino acid residues 105-249) (**Table S1**; **Fig S1**). These data suggest a direct interaction between LegC4 and PA28α, which is conserved in amoebae and upregulated by cytokine signaling in macrophages.

We sought to validate our Y2H data using reciprocal affinity purification, co-immunoprecipitation (IP), and confocal microscopy. We evaluated direct binding between recombinant PA28α and LegC4 by reciprocal affinity chromatography on GST- or His_6_-Myc-tagged fusion proteins produced in *E. coli.* Using glutathione agarose beads, we found that GST-LegC4 retained Myc-PA28α (**Fig 3A**) and GST-PA28α retained Myc-LegC4 (**Fig 3B**). PA28α did not interact with either GST alone or the unrelated *L. pneumophila* effector Lgt1 (**Fig 3A-B**), known to target eukaryotic elongation factor 1A [35,36], which indicates that PA28α does not bind non-specifically to *L. pneumophila* effectors. To determine if PA28α and LegC4 interact in mammalian cells, we co-transfected HEK293T cells to transiently produce epitope-tagged 3xFLAG-LegC4 and either PA28α-Myc or GFP-PA28α fusion proteins. Ectopic expression of PA28α was necessary in these experiments since it is expressed only in response to cytokine signaling [37,38]. Using α-FLAG and α-GFP pull-downs, we found that PA28α-Myc co-immunoprecipitated with 3xFLAG-LegC4 (**Fig 3C**) and that 3xFLAG-LegC4 co-immunoprecipitated with GFP-PA28α (**Fig 3D**), respectively. Together, these data show that LegC4 binds PA28α.

**Figure 3.**
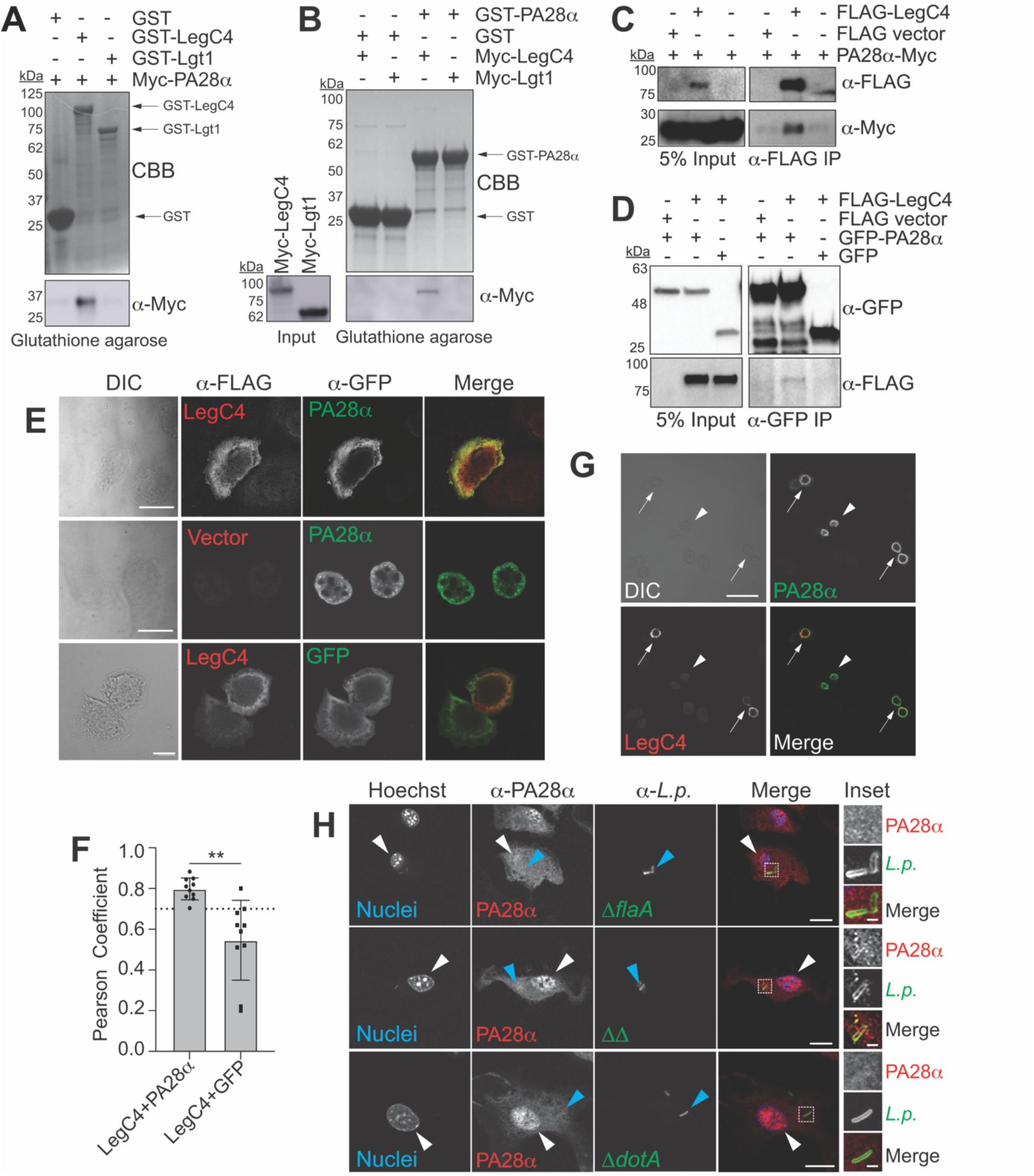
LegC4 interacts with and alters the subcellular localization of PA28α. **(A-B)** Reciprocal affinity chromatography of recombinant GST-fusion proteins from *E. coli* lysates bound to glutathione agarose beads and incubated with lysates from *E. coli* producing His_6_-Myc-fusion proteins. Proteins retained on the beads were eluted and visualized by SDS-PAGE and Coomassie brilliant blue (CBB) stain and α-Myc Western blot. Arrows denote distinct GST-fusion proteins (GST, 26 kDa; GST-LegC4, 113 kDa; GST-Lgt1, 85 kDa; GST-PA28α, 55 kDa; His_6_-Myc-LegC4, 89 kDa; His_6_-Myc-Lgt1, 62 kDa; His_6_-Myc-PA28α, 31 kDa). Data are representative of three experiments. **(C)** Western blot of expression (input) and co-immunoprecipitation of PA28α-Myc with 3xFLAG-LegC4 on a-FLAG conjugated magnetic beads from lysates of co-transfected HEK293T cells. Data are representative of two independent experiments. (**D**) Western blot of expression (input) and co-immunoprecipitation of 3xFLAG-LegC4 with GFP-PA28α on α-GFP-conjugated magnetic beads from lysates of co-transfected HEK 293T cells. Data are representative of two experiments. **(E)** Representative confocal micrographs of transfected HeLa cells producing ectopic 3xFLAG-LegC4 and either GFP-PA28α or GFP stained with α-FLAG and α-GFP antibodies. Scale bar represents 10 µm. **(F)** Pearson Correlation Coefficients (PCC) were calculated for individual cells producing 3xFLAG-LegC4 and either GFP-PA28α or GFP alone (*n*=10) using Fiji ImageJ software. PCC ≥ 0.7 (dashed line) indicates a high confidence association. Asterisks denote statistical significance by Students’ *t*-test (\*\**P*<0.01). **(G)** Confocal micrograph of transfected HeLa cells producing GFP-PA28α (arrowhead) or GFP-PA28α and 3xFLAG-LegC4 (arrows). Scale bar represents 50 µm. **(H)** Confocal micrographs of WT BMDMs infected with *L. pneumophila* strains (MOI of 25) for 7h and stained with α-PA28α (red) and α-*L. pneumophila* (green) antibodies. Cellular nuclei were imaged with Hoechst (blue). White and blue arrowheads indicate BMDM nuclei and intracellular bacteria, respectively. Scale bar represents 10 µm. Inset images are magnified from regions indicated with white boxes. Scale bar represents 1 µm. Images are representative of populations of infected BMDMs over two independent experiments. ΔΔ: Δ*flaA*Δ*legC4*

We further evaluated the biological relevance of LegC4-PA28α binding by evaluating protein localization in cells by confocal microscopy. Ectopically produced 3xFLAG-LegC4 co-localized with GFP-PA28α within transiently transfected HeLa cells since calculated Pearson Correlation Coefficients (PCC) indicated a significant positive correlation between 3xFLAG-LegC4 and GFP-PA28α (Mean PCC = 0.798 ± 0.054) compared to 3xFLAG-LegC4 and GFP (Mean PCC = 0.546 ± 0.196) (**Fig 3E-F**). Interestingly, LegC4 modulated PA28α subcellular localization since GFP-PA28α localized to the nucleus in the absence of LegC4 (**Fig 3E**; **Fig 3G, arrowheads**) but was cytosolically localized within cells co-expressing LegC4 **(Fig 3E; Fig 3G, arrows**) [39]. We subsequently tested whether PA28α localization is influenced by Dot/Icm-translocated LegC4 within *L. pneumophila*-infected BMDMs. WT BMDMs were infected with *L. pneumophila* Δ*flaA*, ΔΔ, or Δ*dotA* for 7.5 h, at which time a replication-competent LCV is established. Fixed cells were permeabilized and stained with α-*L. pneumophila* and α-PA28α antibodies and Hoechst (DNA) and imaged by confocal microscopy. Endogenous PA28α is constitutively produced in immune cells so can be imaged in BMDMs. We found that PA28α was primarily cytosolic within BMDMs infected with *L. pneumophila* Δ*flaA*, but localized robustly to the nucleus of BMDMs infected with either *L. pneumophila* ΔΔ or Δ*dotA* (**Fig 3H, white arrowheads**). Interestingly, we PA28α was present on LCVs harboring *L. pneumophila* ΔΔ but not Δ*flaA* or Δ*dotA* control strains (**Fig 3H, blue arrowheads and inset**). Although altered PA28α localization mediated by translocated LegC4 in naïve infected BMDMs was not sufficient for LegC4 restriction or phagolysosomal fusion in naïve WT BMDMs (Fig 1A, Fig 2B), these data further support a biologically relevant interaction between LegC4 and PA28α, which has not previously identified as a target of bacterial effectors.

### LegC4 restriction phenotypes are suppressed in PA28α-deficient BMDMs

Based on the interaction between PA28α and LegC4, we tested the hypothesis that PA28α contributes to LegC4-mediated restriction of *L. pneumophila*. PA28α-deficient (*Psme1*^-/-^) mice are not commercially available, so we evaluated the role of PA28α using BMDMs derived from commercially-available PA28αβ-deficient (*Psme1/2*^-/-^) mice [5]. PA28α associates with PA28β to form the PA28αβ (11S) proteasome regulator [40]. PA28β is homologous to PA28α but is dependent on PA28α for activity and is hypothesized to enhance the interaction of PA28α with 20S proteolytic proteasome core particles (CPs) [41]. Thus, we evaluated the role of PA28α in LegC4 restriction in BMDMs and *in vivo* using *Psme1/2*^-/-^ mice. We first tested whether loss of PA28αβ relieves the LegC4-mediated growth attenuation within TNF-activated BMDMs. In agreement with our previous work, *L. pneumophila* Δ*flaA* growth was significantly impaired relative to *L. pneumophila* ΔΔ in rTNF-activated WT BMDMs (*P*=0.015); however, growth of *L. pneumophila* Δ*flaA* and ΔΔ did not differ significantly within *Psme1/2*^-/-^ BMDMs (*P*=0.10; **Fig 4A)**. Moreover, overexpression of *legC4* was insufficient to significantly impair *L. pneumophila* replication within naïve *Psme1/2*^-/-^ BMDMs (*P*=0.15) whereas replication was substantially diminished within WT BMDMs (*P*=0.006; **Fig 4B**). Although replication of LegC4-producing bacteria was modestly attenuated in naïve and rTNF *Psme1/2*^-/-^ BMDMs, the lack of statistical significance and differences relative to WT BMDMs provides evidence that PA28αβ activity contributes to LegC4 growth restriction in cultured macrophages.

**Figure 4.**
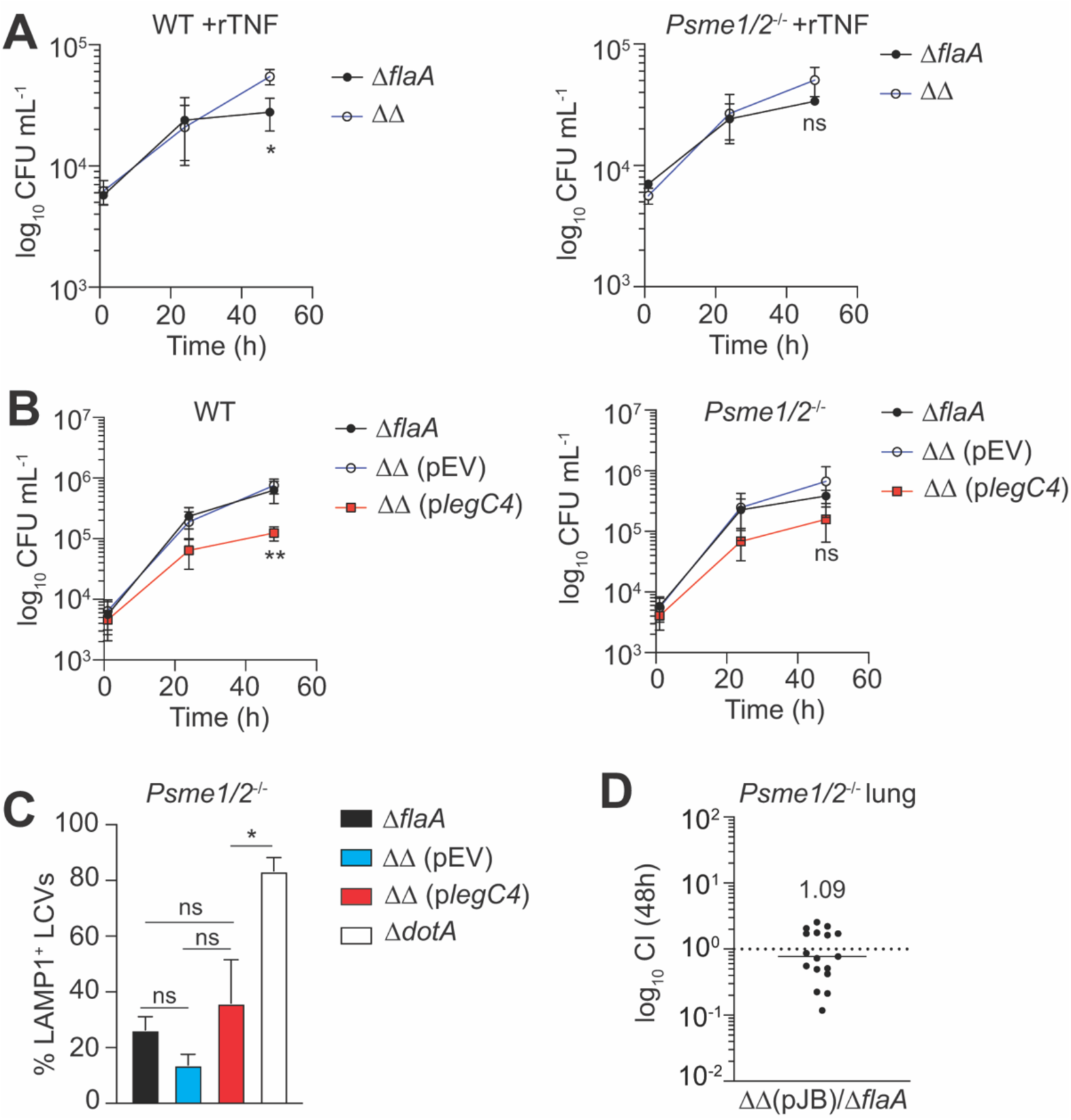
PA28αβ is important for LegC4 restriction phenotypes within cultured BMDMs and the mouse lung *in vivo*. Growth of *L. pneumophila* strains within rTNF-treated **(A)** or naïve **(B)** WT (top) or *Psme1/2*^-/-^ (bottom) BMDMs. Where indicated, BMDMs were treated with 25 ng mL^-1^ rTNF and plasmid expression of *legC4* was induced with 1 mM IPTG. Data shown are mean ± s.d. of pooled results from three independent experiments with triplicate samples per condition in each experiment. Asterisks denote statistical significance by Students’ *t*-test (\*\**P*<0.01; \**P*<0.05). ns, not significant. **(C)** Quantification of LAMP1^+^ LCVs within *Psme1/2*^-/-^ BMDMs in- fected with *L. pneumophila* strains for 9h. Infections were performed in triplicates and LAMP1^+^ LCVs were scored blind (*n*=50 infected cells per coverslip) by immunofluorescence microscopy (*n*=150 per condition per experiment). Data shown are mean ± s.e.m. of pooled results of three independent experiments (*n*=450 total per condition) and asterisks denote statistical significance by two-way ANOVA (\**P*<0.05). ns, not significant. Plasmid expression of *legC4* was induced with 1 mM IPTG. **(D)** Competitive index (CI) values from the lungs of *Psme1/2*^-/-^ mice infected with a 1:1 mixture of *L. pneumophila* ΔΔ (pJB) and *L. pneumophila* Δ*flaA* at 48 h. Each symbol represents an individual animal and data are pooled from three independent experiments. The mean CI value is shown. ΔΔ: Δ*flaA*Δ*legC4*.

We subsequently tested the hypothesis that PA28αβ contributes to LegC4-mediated phagolysosomal fusion with LCVs (Fig 2). Indeed, there were no differences in the abundance of LAMP1^+^ LCVs within *Psme1/2*^-/-^ BMDMs infected with *L. pneumophila* (p*legC4*) compared to *L. pneumophila* Δ*flaA* or ΔΔ (pEV) (**Fig 4C**). PA28ab did not affect phagolysosomal fusion with avirulent *L. pneumophila* Δ*dotA* since significantly more LAMP1+ LCVs were present compared to all other strains (**Fig 4C**). Thus, LegC4-mediated phagolysosomal fusion is abolished in the absence of PA28αβ.

### LegC4 does not confer a fitness disadvantage in the lungs of *Psme1/2*^-/-^ mice

Since loss of PA28αβ was sufficient to suppress LegC4-mediated growth attenuation and phagolysosomal fusion in BMDMs, we hypothesized that LegC4 would not confer a fitness disadvantage on *L. pneumophila* in the lungs of *Psme1/2*^-/-^ mice. We infected *Psme1/2*^-/-^ mice with a 1:1 mixture of *L. pneumophila* ΔΔ (pEV) and Δ*flaA* calculated competitive index values from lung homogenates at 48 h post-infection. Strikingly, *L. pneumophila* fitness was unaffected by LegC4 in the lungs of *Psme1/2*^-/-^ mice (CI=1.09 ± 0.793); **Fig 4D**). In aggregate, our data suggest that PA28αβ is important for LegC4-mediated restriction in cytokine activated BMDMs and the mouse lung *in vivo*.

### The abundance of carbonylated proteins in macrophages is increased in the presence of LegC4 under oxidative stress conditions

PA28αβ is expressed constitutively in cells of the innate immune system and upregulated to maintain the integrity of the proteome under oxidative stress conditions [2,38,40,42]. PA28αβ engages proteolytic proteasome 20S core particles (CPs) to promote rapid ubiquitin-independent proteolysis of oxidant-damaged (carbonylated) proteins [4,43], the abundance of which increases commensurately with the abundance reactive oxygen species (ROS) [44,45]. 20S CPs can degrade carbonylated proteins, but their efficiency is substantially increased when in complex PA28αβ [2,43]. Concomitantly, there is a strong negative correlation between PA28αβ activity and carbonylated protein abundance under oxidative stress conditions [43]. Our Y2H data suggest that LegC4 binds the C-terminal domain of PA28α, which is important for PA28αβ docking on proteasome CPs [46–48]. Thus, we rationalized that carbonylated proteins would be more abundant in the presence of LegC4 under oxidative stress conditions. To directly evaluate LegC4, we generated RAW 264.7 cells stably transfected with 3xFLAG-LegC4 (RAW-LegC4) or empty plasmid vector (RAW-Cntrl). We confirmed 3xFLAG-LegC4 production by confocal microscopy (**Fig 5A**) and Western blot (**Fig 5B**). Immunoprecipitation was necessary to visualize LegC4 since it is produced by macrophages at low abundance. However, the abundance of ectopically produced LegC4 was sufficient to attenuate replication of *L. pneumophila* Δ*flaA* (**Fig 5C**). Attenuated growth was observed at 48 h post-infection, similar to the kinetics of LegC4 restriction within BMDMs (see Fig 1A), which corroborates our microscopy data suggesting that ectopically-produced LegC4 is functional (see Fig 3G-H) and demonstrates that RAW-LegC4 cells are suitable to evaluate LegC4 activity. RAW-LegC4 and -Cntrl cells were incubated in the presence or absence of 10 µM H_2_O_2_ for 24 h and protein carbonyls in cell lysates were quantified by ELISA. We used H_2_O_2_ since it is membrane permeable, stable in solution, and biologically relevant since *L. pneumophila-*infected macrophages in the lung would also be exposed to neutrophil-generated extracellular ROS [26,49]. Indeed, H_2_O_2_-treated RAW-LegC4 cells contained significantly more protein carbonyls than RAW-Cntrl cells (**Fig 5D**). Under native conditions, the concentration of carbonylated proteins was unaffected by LegC4 (**Fig 5D**), which suggests that LegC4 does not itself induce oxidative stress. The abundance of protein carbonyls was unaffected by H_2_O_2_ in RAW-Cntrl cells, likely because upregulation of PA28αβ over the duration of this experiment was sufficient for turnover of damaged proteins.

**Figure 5.**
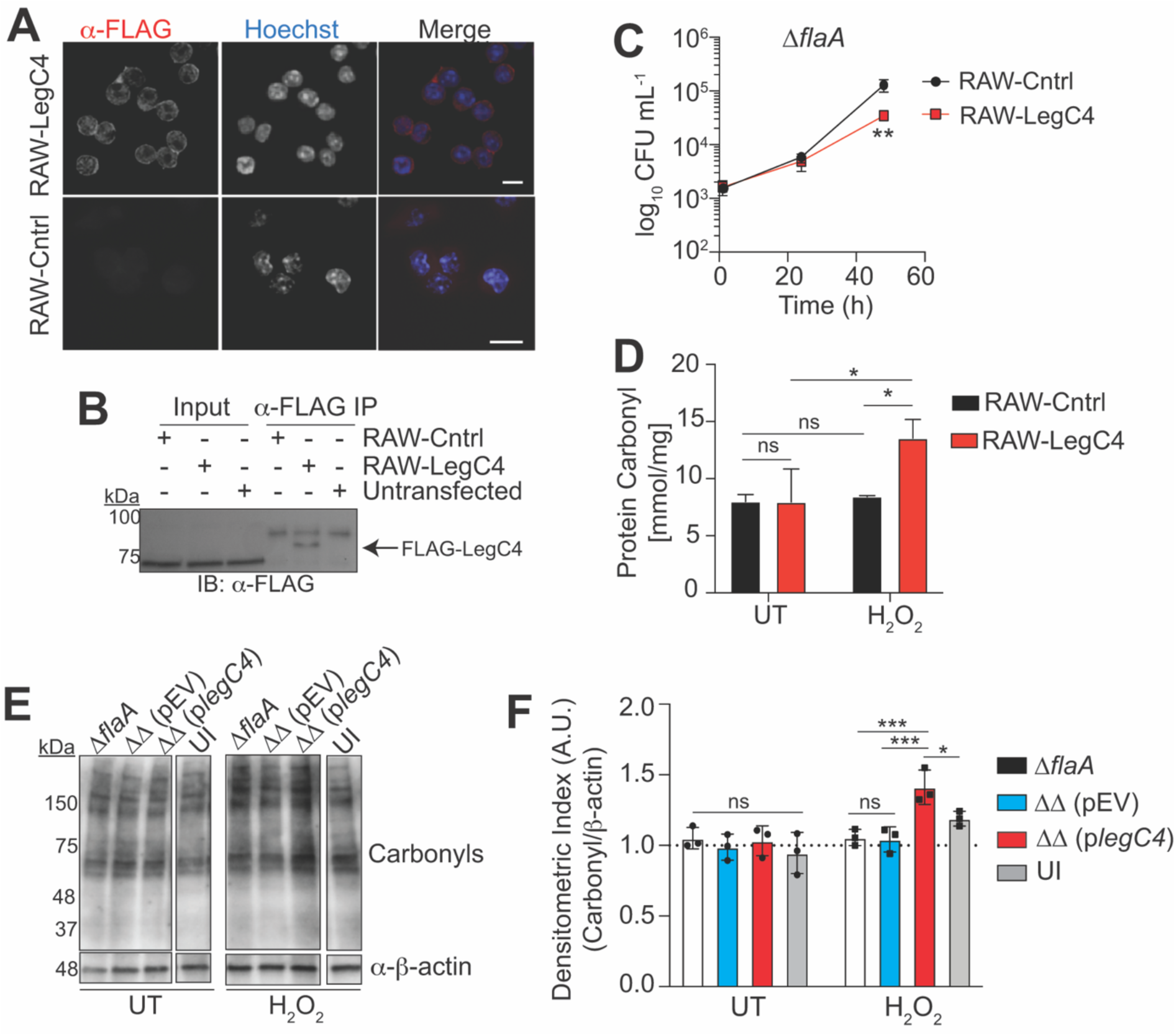
The abundance of carbonylated proteins is increased in the presence of LegC4 under oxidative stress conditions. **(A)** Representative confocal micrographs of RAW-LegC4 and -Cntrl cells stained with α-FLAG antibodies and Hoechst to image nuclei. Scale bar represents 10 µm. **(B)** Western blot of α-FLAG immunoprecipitation from lysates of RAW-LegC4, RAW-Cntrl or untransfected (UT) RAW cells with α-FLAG primary antibody. Arrow indicates 3xFLAG-LegC4 (∼84 kDa). **(C)** Growth of *L. pneumophila* Δ*flaA* within RAW-LegC4 and -Cntrl cells. Data shown are mean ± s.d. of pooled results of three independent experiments with triplicate samples per condition in each experiment. Asterisks denote statistical significance by Students’ *t*-test (\*\**P*<0.01). **(D)** Quantification of protein carbonyls by ELISA in lysates of RAW-LegC4 and -Cntrl cells incubated in low-serum media for 24 h in the presence or absence of 10 µM H_2_O_2_. Data shown are mean ± s.e.m. and representative of two independent experiments. Asterisks denote statistical significance by two-way ANOVA (\**P*<0.05; ns, not significant). UT: untreated. **(E)** Western blot for protein carbonyls and β-actin (loading control) in lysates of *Tnfr1*^-/-^ BMDMs infected for 4 h with the indicated strains (MOI of 50) in the presence or absence of 50 µM H_2_O_2_. Uninfected (UI) samples are from the same experiment but shown separately to remove irrelevant lanes. ΔΔ: Δ*flaA*Δ*legC4*; plasmid expression of *legC4* was induced with 1 mM IPTG. Data are representative of three independent experiments. **(F)** Densitometric quantification of carbonylated proteins from Western blots. Signal intensity from protein carbonyl Western blots was normalized to the β-actin loading control for each sample and averaged ratios were normalized to Δ*flaA*-infected cells. Data shown are mean ± s.d. of values from three independent experiments. Asterisks denote statistical significance by two-way ANOVA (\*\**P*<0.01; \**P*<0.05). ns, not significant.

**Figure 6.**
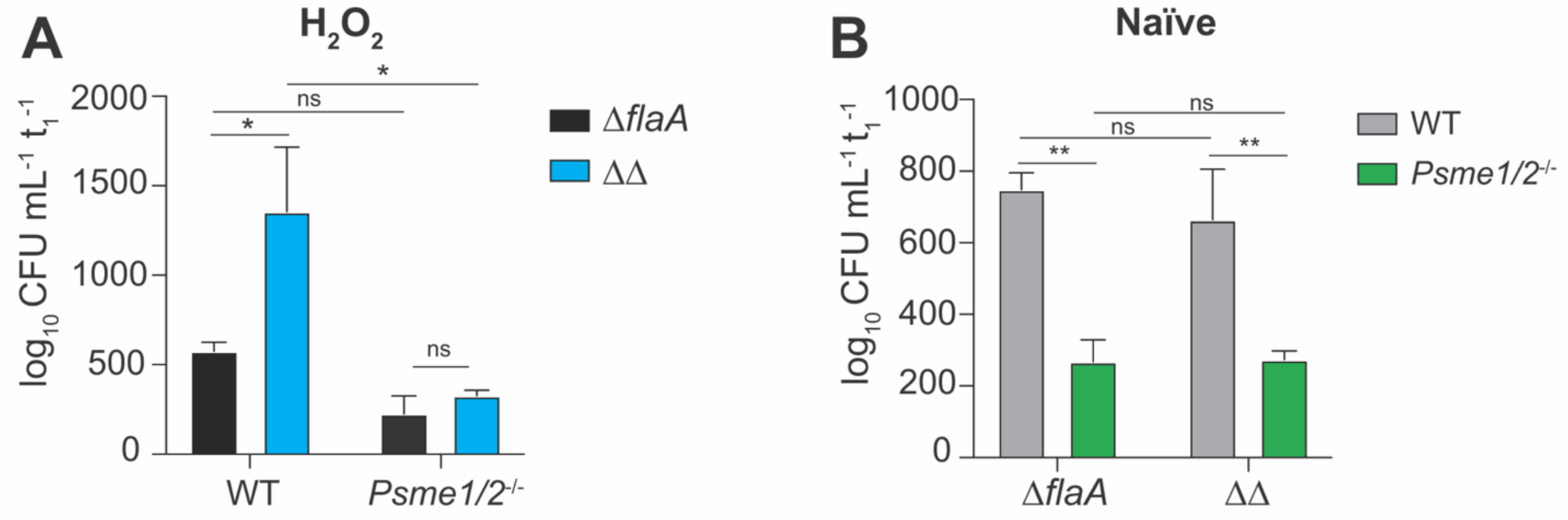
LegC4 restriction occurs under oxidative stress conditions and is phenocopied by loss of PA28αβ. Normalized CFU recovered from **(A)** 20 µM H_2_O_2_-treated or **(B)** naïve WT or *Psme1/2*^-/-^ BMDMs infected for 48 h with either *L. pneumophila* Δ*flaA* or ΔΔ. CFU of each strain recovered at 48 h was normalized to internalized bacteria at 1h post-infection for each condition. Data shown are mean ± s.e.m. of pooled results from three independent experiments with triplicate samples per condition in each experiment. Asterisks denote statistical significance by two-way ANOVA (\*\**P*<0.01, \**P*<0.05). ns, not significant.

We subsequently tested whether LegC4 increases the abundance of protein carbonyls within *L. pneumophila*-infected BMDMs. *Tnfr1*^-/-^ BMDMs were used to control for any LegC4-mediated differences in TNF secretion [20] and, consequently, intracellular ROS production [50]. *Tnfr1*^-/-^ BMDMs were infected with *L. pneumophila* Δ*flaA*, ΔΔ (pEV), or ΔΔ (p*legC4*) for 4 h in the presence or absence of 50 µM H_2_O_2_ and protein carbonyls were quantified by Western blot and densitometry. We found that carbonylated proteins were more abundant within BMDMs infected with *L. pneumophila* ΔΔ (p*legC4*) compared to *L. pneumophila* Δ*flaA* and ΔΔ (pEV)-infected and uninfected BMDMs when oxidative stress was induced with H_2_O_2_ (**Fig 5E-F**). The abundance of protein carbonyls was not affected by LegC4 within naïve untreated BMDMs, which further suggests that LegC4 itself does not induce oxidative stress. Protein carbonyl abundance was not affected by either LegC4 or *L. pneumophila* infection within untreated (UT) BMDMs (**Fig 5E-F**), likely because the amount ROS produced in response to *L. pneumophila* infection under these conditions is insufficient for detectable differences in protein carbonyls by Western blot. Together, these data support our hypothesis that LegC4 perturbs proteostasis in macrophages under oxidative stress conditions and suggest that LegC4 may function by impairing PA28αβ activity.

### Oxidative stress is sufficient for LegC4 restriction in PA28αβ-producing BMDMs

Our current data suggest role(s) for PA28αβ and oxidative stress in LegC4 restriction; thus, we hypothesized that oxidative stress would be sufficient for LegC4-mediated restriction within BMDMs producing PA28αβ. To test this, we quantified replication of *L. pneumophila* Δ*flaA* and ΔΔ within WT and *Psme1/2*^-/-^ BMDMs treated with 20 µM H_2_O_2_ to induce oxidative stress. Indeed, oxidative stress was sufficient for LegC4 restriction since *L. pneumophila* ΔΔ replicated to significantly higher levels than *L. pneumophila* Δ*flaA* in WT BMDMs (**Fig 6A**), showing that oxidative stress is sufficient for LegC4 restriction. Importantly, the growth advantage associated with Δ*legC4* mutation was abolished in PA28αβ-deficient BMDMs, but replication of LegC4-producing bacteria was unaffected (**Fig 6A**). These data show that LegC4 impairs *L. pneumophila* replication under oxidative stress conditions via a mechanism involving PA28αβ.

### *L. pneumophila* replication is impaired in *Psme1/2*^-/-^ BMDMs irrespective of LegC4

Our data suggest that LegC4 restriction is mediated by subversion of PA28αβ. Thus, it’s possible that the dose-dependency of LegC4 in naïve BMDMs results from stoichiometric differences in its abundance relative to PA28αβ. Thus, we rationalized that *L. pneumophila* replication would be suppressed in naïve *Psme1/2*^-/-^ BMDMs irrespective of LegC4 and found that, indeed, both *L. pneumophila* Δ*flaA* and ΔΔ strains were attenuated for replication within *Psme1/2*^-/-^ BMDMs (**Fig 6B**).

Importantly, impaired replication in *Psme1/2*^-/-^ BMDMs was independent of LegC4 since *L. pneumophila* Δ*flaA* and ΔΔ were equally attenuated (**Fig 6B**). These data show that PA28αβ-deficient macrophages are more restrictive restriction to *L. pneumophila* and suggest a novel role for PA28αβ in host defense against intracellular pathogens in macrophages.

### Neither TNF production nor cytotoxicity are potentiated by loss of PA82αβ in *L. pneumophila*-infected BMDMs

Our data suggest that LegC4 enhances phagolysosomal fusion with LCVs via subversion of PA28αβ activity and that loss of PA28αβ is sufficient to impair L. pneumophila intracellular replication. However, impaired bacterial growth may also result from increased TNF production or death of infected BMDMs. We quantified TNF in the supernatants of WT and *Psme1/2*^-/-^ BMDMs infected with *L. pneumophila* Δ*flaA*, ΔΔ (pEV), ΔΔ (p*legC4*) and Δ*dotA* (negative control). At 8h post-infection, there was a modest but statistically significant decrease in TNF secretion from *Psme1/2*^-/-^ BMDMs infected with *L. pneumophila* ΔΔ (p*legC4*) (*P*=0.049) but no other PA28αβ-mediated differences were observed (**Fig S2A**). At 24 h post-infection, significantly less TNF was present in supernatants of *Psme1/2^-^*^/-^ BMDMs infected with *L. pneumophila* Δ*flaA* or ΔΔ (pEV) compared to WT BMDMs but no differences in secretion from BMDMs infected with *L. pneumophila* ΔΔ (p*legC4*) (**Fig S2B**). Furthermore, viability of *L. pneumophila*-infected was not affected by either LegC4 or PA28αβ since there were no differences in lactate dehydrogenase (LDH) activity in culture supernatants (**Fig S3**). These data suggest neither LegC4 nor PA28αβ restriction phenotypes result from increased TNF production or death of infected BMDMs.

### Inflammasome activation is dispensable for LegC4-mediated restriction

Our data suggest that LegC4 perturbs proteostasis and potentiates oxidative stress. Sustained oxidative and proteotoxic stress and proteostasis perturbations can activate the NLRP3 inflammasome, which culminates in activation of caspase-1, an effector caspase responsible for pyroptotic cell death and inflammatory cytokine secretion. However, pharmacological inhibition of the NLRP3 inflammasome with MCC950 did not affect LegC4 restriction since *L. pneumophila* ΔΔ (p*legC4*) was still attenuated for replication compared to *L. pneumophila* ΔΔ (pEV) and Δ*flaA* within WT BMDMs (**Fig S4A**). We also found that LegC4 restriction was preserved within BMDMs derived from caspase-1-deficient (*Casp1*^-/-^) mice (**Fig S4B**). Within cultured macrophages, pyroptotic cell death is responsible for *L. pneumophila* restriction downstream of inflammasome activation. Our data suggest that LegC4 does not enhance LDH release from infected BMDMs (**Fig S3**); however, pyroptotic cell death is more pronounced within BMDMs primed with toll-like (TLR) receptor agonists [51,52]. Since TLR2 is important for host defense against *L. pneumophila* [25], we evaluated LDH release from *L. pneumophila*-infected BMDMs pretreated with the TLR2 agonist PAM_3_CSK_4_. Similar to our previous results (**Fig S3**), LDH release from BMDMs primed with PAM_3_CSK_4_ was unaffected by LegC4 (**Fig S4C**). These data suggest that LegC4-mediated restriction occurs independently of NLRP3- and caspase-1-dependent pyroptotic cell death and support an overarching model whereby subversion of PA28αβ by LegC4 augments lysosomal pathogen killing under inflammatory and oxidative stress conditions.

## Discussion

The Dot/Icm effector LegC4 is *bona fide L. pneumophila* virulence factor in a natural host amoeba but confers a dose-dependent fitness disadvantage on *L. pneumophila* in cytokine-activated macrophages and in a mouse model of Legionnaires’ Disease [20,21]. LegC4 restriction represents a unique mode of effector-mediated immunity since its activity augments existing inflammatory responses in macrophages. In this study, we have uncovered mechanistic insights into LegC4 restriction and a new role for PA28αβ in cell-autonomous host defense against *L. pneumophila*. We propose a model whereby subversion of host PA28αβ by LegC4 perturbs proteostasis under oxidative stress conditions, resulting in increased lysosome biogenesis and phagolysosomal fusion with LCVs (**Fig S5**). This model is supported by our data showing that (1) LegC4 enhances cytokine-mediated phagolysosomal fusion with LCVs; (2) LegC4 binds and modulates the subcellular localization of PA28α; (2) PA28αβ is required for the effects of LegC4 in BMDMs and *in vivo*; (3) PA28αβ substrates are more abundant in the presence of LegC4; (4) oxidative stress is sufficient for LegC4 restriction; and (5) loss of PA28αβ was restrictive to *L. pneumophila* irrespective of LegC4. This work is the first to identify PA28α as a target for a bacterial effector protein and a role for PA28αβ in innate and effector-mediated host defense against an intracellular bacterial pathogen.

*L. pneumophila* is an accidental pathogen of humans that rarely transmits between infected individuals. Thus, an evolutionary basis for the LegC4-PA28α interaction can be explained by LegC4 targeting the PA28α homolog in amoebae. Indeed, the natural host *A. castellanii*, in which loss of *legC4* confers a growth defect [21], encodes a PA28α homolog (*Ac*PA28) that shares 34% identity with mouse PA28α. Sequence identity between PA28α and *Ac*PA28 is primarily in the PA28α C-terminal domain, which is important for engaging 20S CPs and was contained in all PA28α clones identified in our Y2H screen (**Table S1**). Since PA28αβ substrates were more abundant in the presence of LegC4 (**Fig 5**), we postulate that LegC4 binding impairs PA28-CP complex formation. The function of *Ac*PA28 has not been described; however, it is likely a proteasome regulator based on structural studies showing that mammalian PA28α and evolutionary distant PA28s are similar and all associate with 20S CPs [47,48]. The benefit of inhibiting PA28 in the natural host is unclear, but sequestering PA28 from CPs may preserve the integrity of the constitutive 26S proteasome, a central player in *L. pneumophila*’s virulence strategy [53]. We are currently investigating the molecular mechanism by which LegC4 modulates PA28 activity and its role in *L. pneumophila* virulence in natural host amoebae.

Virulent *L. pneumophila* establish a replicative LCV by avoiding endosomal maturation of their phagosome. The current dogma dictates that lysosomal fusion with LCVs at early time points during infection (≤ 16 h) and *L. pneumophila* intracellular replication are mutually exclusive [33,54]. We found that plasmid expression of *legC4* increased the abundance of LAMP1 on LCVs in within WT BMDMs, similar to what is seen for the avirulent Dot/Icm-deficient strain. LegC4-mediated phagolysosomal fusion was suppressed in *Tnfr1*^-/-^ BMDMs, but significantly more LAMP1+ LCVs were present relative to cells infected with *L. pneumophila* Δ*flaA* and ΔΔ (pEV) control strains. This is intriguing since plasmid expression of *legC4* does not affect *L. pneumophila* replication within *Tnfr1*^-/-^ BMDMs [20] (**Fig 1D**). Based on our model that phagolysosomal fusion results from oxidative stress (**Fig S5**), we postulate that the TNFR1-independent increase in LAMP1+ LCVs within in cells infected with the *legC4* overexpressing strain may result from ROS produced in response to phagocytosis or pattern-recognition receptor signaling [55,56]. A potential explanation for this is that LegC4 may target other host proteins when PA28α is absent, such Ecm29 or PA28γ, which were identified in our Y2H screen (**Table S1**). Ecm29 is a particularly interesting candidate since it disassembles 26S proteasomes under oxidative stress conditions to liberate free 20S CPs that can then associate with PA28αβ [57–59]. This may also explain the modest and non-significant LegC4-mediated growth attenuation retained in *Psme1/2*^-/-^ BMDMs (**Fig 4A-B**). Interestingly, robust phagolysosomal fusion with LCVs was not observed in *Psme1/2*^-/-^ BMDMs although these cells are restrictive to *L. pneumophila* (**Fig 6B**). A potential explanation for this observation is that less TNF is produced by *L. pneumophila* ΔΔ (p*legC4*)-infected *Psme1/2*^-/-^ BMDMs compared to WT BMDMs (**Fig S2**), although this is unlikely to account for loss of LegC4 growth restriction since phenotypes are apparent in rTNF-treated BMDMs (**Fig 4A**). A mechanistic connection between PA28αβ and phagolysosomal fusion can be explained by a compensatory upregulation of lysosome biogenesis and degradation pathways resulting from impaired proteasome activity and oxidative stress [3,60,61]. Our data suggest that lysosomes are central to LegC4 restriction, but the underlying mechanism(s) of phagolysosomal fusion with LCVs and bacterial killing remain to be fully elucidated.

Lysosomal degradation pathways, including autophagy, are cytoprotective under inflammatory and oxidative stress conditions and central to cytokine-mediated cell-autonomous pathogen restriction. Lysosomal restriction within IFN-γ-activated macrophages involves immunity-related GTPases (IRGs), which localize to and facilitate lysosomal trafficking to pathogen-containing phagosomes [62]. Although IRGs may contribute to restriction of LegC4-producing *L. pneumophila* strains within IFN-γ-activated macrophages [63], IFNGR1 signaling was dispensable for LegC4 restriction (Fig 1E-F), which suggests that IRGs are not central drivers of LegC4-mediated *L. pneumophila* restriction. It’s tempting to speculate that autophagy contributes to LegC4 restriction since autophagy genes are upregulated to clear protein aggregates and protect against TNF-induced death in myeloid cells [64]. Autophagy also prevents aberrant inflammasome-mediated pyroptotic cell death below a certain threshold of inflammasome agonist [65]. The NLRC4 inflammasome is dispensable for LegC4-mediated restriction [20,21] and we also found no role for the NLRP3 inflammasome (**Fig S4**), which is activated by both oxidative and proteotoxic stress [66]. Whether autophagy contributes to LegC4-mediated restriction is unknown and warrants investigation since several *L. pneumophila* effectors impair global autophagy in host cells [67–69].

Using several complementary approaches, we found that LegC4 binds and PA28α and influences its localization in transfected and infected cells. The change in PA28α localization supports an interaction with LegC4, but the underlying mechanism by which it occurs is unclear. We were unable to detect Dot/Icm-translocated LegC4 by confocal microscopy within infected BMDMs, likely because the abundance is below the limit of detection, which is a common challenge in the study of bacterial effectors that do not localize to organellar membranes [70,71]. Our data suggest that the interaction between LegC4 and PA28α may drive localization changes in transfected cells; however, the mechanism is likely more complex since presumed stoichiometric imbalances between PA28α and Dot/Icm-translocated LegC4 make it unlikely that binding is sufficient. Thus, in infected macrophages, LegC4-mediated changes in PA28α localization may result from a need for additional nuclear PA28αβ to clear proteins damaged by the oxidative burst in the presence of LegC4. However, the extent of oxidative damage under these conditions is clearly insufficient to impair *L. pneumophila* replication since endogenous LegC4 does not affect bacterial replication within naïve BMDMs. Interestingly, we also found endogenous PA28α on LCVs harboring LegC4-deficient *L. pneumophila*, which raises the possibilities that LegC4 binding or activity sequesters PA28α in the cytosol away from proteasome complexes associated with the LCV [63,72]. Although changes in PA28α localization support its functional modulation by LegC4, the molecular underpinnings and consequences of LegC4-PA28α binding, especially based on stoichiometric differences in protein abundance during infection, on PA28ab activity and pathogen restriction require further investigation.

A role for PA28αβ and oxidative stress in LegC4 restriction was also revealed by the sufficiency of hydrogen peroxide (H_2_O_2_) for impaired growth in WT BMDMs. H_2_O_2_ is routinely used to induce oxidative stress in laboratory models since both superoxide (O_2_^•-^) and hydroxyl (OH^•^) radicals are unstable in solution. Within immune cells, phagocytosis, pro-inflammatory cytokines, and PRR signaling increase production of ROS by phagocyte NADPH oxidase (NOX2) and mitochondria [50,55,56,73–75]. Intracellular ROS contribute to cell-autonomous pathogen restriction but ROS are also released into the extracellular space via plasma membrane-bound NOX2 or by passive diffusion across membranes (H_2_O_2_) [49,76]. Interestingly, ROS play a negligible role in host defense against *L. pneumophila* in cultured naïve macrophages, likely owing NOX2 exclusion from LCVs [26,77,78]. However, in the lung, ROS are produced abundantly by neutrophils and neutrophil ROS is important for host defense against *L. pneumophila* [26]. Thus, infected alveolar macrophages are exposed to both extracellular ROS produced by lung neutrophils and infection- and cytokine-induced intracellular ROS [49,79,80]. When we treated BMDMs with exogenous H_2_O_2_ to recapitulate the *in vivo* environment, endogenous LegC4 was sufficient to impair *L. pneumophila* replication (**Fig 6A**); thus, the fitness disadvantage associated with LegC4 *in vivo* may be due to oxidative stress. Furthermore, it is tempting to speculate that the dose-dependency of LegC4 restriction may correlate with the abundance of ROS. ROS are microbicidal, but this activity is unlikely to be involved in LegC4 restriction since (1) *L. pneumophila* has multiple complimentary strategies to protect against oxidative stress and can withstand up to 2 mM H_2_O_2_ *in vitro* (orders of magnitude greater than intracellular concentrations) [78,81–83]; and (2) replication of LegC4-deficient *L. pneumophila* was unaffected by H_2_O_2_ treatment (**Fig 6A-B**). While our data do not preclude roles for additional host factors or cytokine-driven cellular processes in LegC4 restriction of *L. pneumophila*, they support a central role for oxidative stress.

Our data suggest that oxidative stress is sustained in the presence of LegC4 since oxidatively damaged (carbonylated) proteins were more abundant. Carbonylated proteins are used as a biomarker for oxidative stress since proteins and nascent polypeptides are highly susceptible to oxidative damage [44,45]. Importantly, increased protein carbonyl abundance is also indicative of the inability of cellular proteasomes and chaperones to keep pace with the rate of damage [84]. Thus, it is conceivable that impaired PA28αβ activity is indeed responsible for increased protein carbonyl abundance in the presence of LegC4. Nascent polypeptides and misfolded proteins are more susceptible to oxidative damage [85]; thus, impaired co-translational chaperone activity, such as inactivation of Hsp70 by the effector LegK4 [86], may also contribute to carbonylated protein abundance in *L. pneumophila*-infected macrophages. Accumulation of heavily oxidized polypeptides results in formation of aggregates, which are impervious to proteasomal degradation and must be cleared by lysosomal degradation pathways. We think that a global increase in carbonylated proteins drives increased phagolysosomal activity in the presence of LegC4; however, all proteins are not equally susceptible to oxidative damage and individual oxidized species may be involved. For example, the actin cytoskeleton, which plays a critical role in vesicular transport, is deeply influenced by cellular redox status [87]. Filamentous (F)-actin is involved in phagolysosomal fusion with intracellular *L. pneumophila* and pathogen restriction downstream of the caspase-11 inflammasome [88–91]. Caspase-11 promotes phagolysosomal fusion with intracellular pathogens by mediating phosphorylation and inactivation of the actin depolymerization factor cofilin [88,91]. Phosphorylated cofilin is highly susceptible to oxidative damage, and its oxidation may therefore enhance phagolysosomal fusion with LCVs [92].

In summary, we have uncovered a novel mechanism of effector-mediated host defense and a new role for PA28αβ and oxidative stress in cell-autonomous pathogen restriction. Interestingly, PA28αβ is targeted by several viral pathogens, likely as a means to suppress antigen presentation on MHC class I [93–95]. Impaired antigen presentation is unlikely to play a role in *L. pneumophila* restriction since phenotypes are observed in cultured macrophages and during acute lung infection, prior to initiation of an adaptive immune response. This work is the first to show bacterial pathogen targeting of PA28α and suggest that suppression of PA28αβ activity and aberrant proteostasis may be a pathogen-triggered host defense mechanism to potentiate existing inflammatory responses. The ability to enhance the antimicrobial potential of activated macrophages has exciting implications for development of host-centric immunotherapeutics and future studies will provide further insight into the ubiquity and mechanism(s) of pathogen restriction induced by proteotoxic stress and suppression of PA28αβ activity.

## Materials & Methods

### Bacterial Strains, plasmids, oligonucleotide primers, and reagents

*L. pneumophila* strains, plasmids, and primers used in this study are listed in **Table S2, Table S3,** and **Table S4,** respectively. *L. pneumophila* strains were cultured on supplemented charcoal N-(2-Acetamido)-2-aminoethanesulfonic acid (ACES)-buffered yeast extract (CYE) and grown at 37°C, as described [96]. Liquid cultures were grown overnight with shaking at 37°C in supplemented ACES-buffered yeast extract (AYE) medium, as described [20,97]. CYE was supplemented with 10 µg mL^-1^ chloramphenicol for plasmid maintenance and *legC4* expression was induced with 1 mM isopropyl-β-D-1-thiogalactopyranoside (IPTG; GoldBio) where indicated. *Escherichia coli* strains used for cloning (Top10; Invitrogen) and protein expression [BL21 (DE3); New England Biolabs] were grown at 37 °C in Luria-Bertani (LB) medium supplemented with antibiotics as appropriate for plasmid selection [50 µg mL^-1^ kanamycin (GoldBio), 25 µg mL^-1^ chloramphenicol (GoldBio), or 100 µg mL^-1^ ampicillin (GoldBio)]. Protein expression in BL21 (DE3) was induced with 1 mM IPTG. Unless otherwise indicated, all chemicals were obtained from Millipore Sigma (St. Louis, MO).

### Mice and bone marrow-derived macrophages

Mice on C57BL/6 background were purchased from the Jackson Laboratories (Bar Harbor, Maine). Wild-type, *Tnfr1*^-/-^, *Ifngr1*^-/-^, and *Myd88*^-/-^ mice were purchased from live repositories and *Psme1/2*^-/-^ mice were recovered from cryopreserved embryos and bred for homozygotes at the Jackson Laboratories (Bar Harbor, ME). In-house colonies were maintained in specific pathogen-free conditions at Kansas State University and Michigan State University.

Bone marrow was harvested from seven-to twelve-week-old mice as described [98]. BMDMs were generated by differentiation in RPMI supplemented with 20% heat-inactivated fetal bovine serum (HI-FBS) (Biowest, Riverside, MO) and 15% L929 cell supernatant for 6 days prior to seeding for infection. Femurs from C57Bl/6 *Casp1*^-/-^ mice were a gift from Dr. Russell Vance (University of California, Berkeley).

All experiments involving animals were approved by the Kansas State University and Michigan State University Institutional Animal Care and Use Committees (KSU Protocols: 4022, 4501, and 4386; MSU protocol: PROTO202200365) and performed in compliance with the Animal Welfare Act and NIH guidelines. Animals were euthanized in accordance with American Veterinary Medical Association Guidelines.

### Competitive Index

Six-to twelve-week-old sex- and age-matched mice were anesthetized with ketamine/xylazine and infected via the intranasal route as previously described [20]. Mixed bacterial inoculums (1:1) containing a total of 5 x 10^6^ bacteria were diluted and plated on selective medium (10 µg mL^-1^ chloramphenicol for plasmid selection). At 48 h post-infection, mice were euthanized and homogenates from extracted lung tissue were plated on selective media, as described [20]. CFU were enumerated and used to calculate CI values [(CFUcm^R^_48h_/CFUwt_48h_)/(CFUcm^R^_IN_)/CFUwt_IN_)].

### Molecular Cloning

Plasmids were generated for stable and transient ectopic production in mammalian cells and recombinant protein production in *E. coli*. For production of 3xFLAG-LegC4, *legC4* was amplified from *L. pneumophila* genomic (g)DNA using LegC4BamHI-F/LegC4NotI-R primer pairs (**Table S4**) and cloned as a BamHI/NotI fragment into 3xFLAG 4/TO (**Table S3**) [99]. To produce GFP-PA28α, *psme1* was amplified from pCMV-3Tag-4a::*psme1* (purchased from Genscript, Piscataway, New Jersey) using Psme1Sal1-F/Psme1BamHI-R primer pairs (**Table S4**) and cloned as a Sal1/BamH1 fragment into pEGFPC1 (Clontech). For production of GST- and His_6_-Myc-fusion proteins, open reading frames were cloned into pGEX-6P-1 and pT7HMT vectors, respectively. *psme1* was PCR-amplified from pCMV-3Tag-4a::*psme1* and *legC4* PCR-amplified from *L. pneumophila* genomic (g)DNA using primers Psme1-BamHI-F/Psme1-NotI-R and LegC4-BamHI-F3/LegC4-NotI-R, respectively (**Table S4**). Each of the amplified fragments were cloned as a BamHI/NotI fragment into either pGEX-GP-1 or pT7HMT. pGEX::*lgt1* and pT7HMT::*lgt1* were constructed previously [100]. Confirmation of DNA sequences was done by Sanger sequencing (Eton Biosciences Inc.). Plasmid DNA was transformed into chemically competent *E. coli* Top10 (Invitrogen) or BL21 (DE3) (New England Biolabs) for plasmid purification and recombinant protein production, respectively. Plasmid DNA was purified for transfection into mammalian cell lines using a ZymoPURE II Plasmid Maxiprep Kit (Zymo Research).

### Recombinant protein production and affinity chromatography

*E. coli* BL21 (DE3) overnight cultures were sub-cultured into fresh LB medium containing the appropriate antibiotics at 1:100 and grown with shaking for 3h at 37°C. Recombinant protein production was induced with 1mM IPTG and cultures were grown overnight with shaking at 16°C. Bacteria were harvested by centrifugation at 1,400 r.c.f. for 10 min at 4°C, washed with ice-cold PBS and incubated in bacterial lysis buffer [50mM Tris pH 8.0, 100mM NaCl, 1mM EDTA, 200µg mL^-1^ lysozyme, and complete protease inhibitor (Roche)] for 30 min on ice followed by addition of 2 mM DTT. Bacteria were sonicated and lysates were clarified by centrifugation at 21,000 r.c.f. for 15 min at 4°C, as described [100]. Clarified lysates harboring GST-fusion proteins were incubated with magnetic glutathione agarose beads (Pierce) for 1hr at 4°C with rotation and washed twice in Glutathione wash buffer [125mM Tris pH 7.4, 250mM NaCl, 1mM DTT, 1mM EDTA]. Clarified lysates harboring His_6_-Myc-fusion proteins were added to the beads and incubated at 4°C with rotation for 1hr. Beads were washed twice with Glutathione wash buffer and proteins were eluted by boiling in Laemmli sample buffer. Proteins were separated by SDS-PAGE followed by Coomassie brilliant blue staining and Western blot (see below).

### Cell Culture and Transfections

HEK 293T, HeLa, and RAW 264.7 cells (gifts from Dr. Craig Roy, Yale University) were maintained at 37°C and 5% CO_2_ in Dulbecco’s Modified Eagle Medium (DMEM) supplemented with 10% HI-FBS. THP-1 cells were purchased from the American Type Culture Collection (ATCC) and maintained in RPMI supplemented with 10% HIFBS and differentiated with 100 nM phorbal 12-myristate 13-acetate (PMA) for 4 days prior to infection with *L. pneumophila* (see below). All cell lines were used between passage 4-20.

HEK 293T cells were transfected with purified plasmid DNA (**Table S2**) for 24 h using calcium phosphate as described [101]. HeLa cells were transfected for 24 h using jetPRIME transfection reagent according to manufacturer’s guidelines. Two hours before transfection, HeLa cells were washed and incubated in low-serum media (DMEM/4% HIFBS). Media were replaced with DMEM 10% HIFBS four hours post-transfection and assayed 24 h post-transfection.

To generate RAW 264.7 cells with stable plasmid integrations, cells were transfected with pcDNA::*3xflag-legC4* or pcDNA::*3xflag* vector (**Table S2**) using a Nucleofector 2b electroporator (Lonza, Basel, Switzerland) and cultured over 14 days with Zeocin selection (200 -1000 µg mL^-1^). 3xFLAG-LegC4 expression was confirmed by Western blot and confocal microscopy using a-FLAG antibodies (see below) and stable RAW 264.7 cells were maintained in culture in the presence of 200 µg mL^-1^ Zeocin. For *L. pneumophila* infections, cells were seeded one day prior to infection and infected in the absence of Zeocin.

### Western blot

Boiled protein samples were separated by SDS-PAGE and transferred to polyvinylidene difluoride (PVDF) membrane using a BioRad TransBlot semidry transfer apparatus. Membranes were incubated in blocking buffer [5% non-fat milk powder dissolved in Tris-buffered saline-0.1% Tween-20 (TBST)]. Primary antibodies [rabbit α-FLAG (Sigma-Aldrich, F1804), rabbit α-Myc (Cell Signaling, 2278S), rabbit α-GFP (Abcam, ab6556)] were used at 1:1,000 in blocking buffer and detected with horseradish peroxidase (HRP)-conjugated secondary antibodies (1:5,000; ThermoFisher). Membranes were washed in TBST followed by addition of enhanced chemiluminescent (ECL) reagent (GE Amersham) and visualization using an Azure c300 Dark-room Replacer or a GE Amersham Imager 600.

### Immunoprecipitation

Transiently transfected HEK 293T cells or RAW 264.7 cells were washed with phosphate-buffered saline and lysed in ice-cold NP-40 buffer [1% non-iodet P40 (v/v), 20 mM Tris pH 7.5, 150 mM NaCl, 10 mM Na_4_P_2_O_7_, 50 mM NaF, complete protease inhibitor (Roche)]. Lysates were clarified and proteins were immunoprecipitated using α-FLAG M2 magnetic beads or protein G-conjugated Dynabeads (ThermoFisher) pre-incubated with rabbit α-GFP antibody (Abcam, ab6556), as indicated and according to manufacturer’s instructions. Input samples were collected from cell lysates prior to incubation with beads. Beads and input samples were resuspended in 3x Laemmli sample buffer for SDS-PAGE and Western blot analysis.

### Protein carbonyl assays

Protein carbonyls were quantified using the OxiSelect Protein Carbonyl ELISA Kit (Cell Biolabs) following manufacturer’s instructions. Briefly, RAW 264.7 cells were seeded at 2x10^6^ in a 10 cm tissue culture dishes in DMEM 10% HIFBS. The next day, media were aspirated, and cells were incubated in DMEM/2.5% HIFBS in the presence or absence of 10 µM H_2_O_2_. After 24 h, cells were washed twice in PBS and lysed in ice-cold 500 µL of detergent-free lysis buffer [25 mM HEPES pH 7.4, 150 mM NaCl, 10 mM MgCl_2_, 1 mM EDTA, 2% glycerol (v/v)] supplemented with complete protease inhibitor cocktail (Roche). Cell lysates were scraped into a pre-chilled micro-centrifuge tube and incubated with agitation for 30 min at 4°C followed by sonication at 40% intensity three times in 10 second intervals. Lysates were clarified by centrifugation at 4°C for 20 min at 13,000 r.c.f. Supernatants were transferred to a fresh tube, snap frozen in liquid nitrogen for 2 min and stored at -80°C until use. Protein concentrations were quantified using a Coomassie Plus (Bradford) Protein Assay (Pierce) and diluted to 10 µg mL^-1^. Samples and standards were adsorbed to a 96-well protein binding plate and ELISA was performed according to manufacturer’s instructions. Absorbance at 450 nm was quantified on a BioTek Epoch2 microplate reader.

Protein carbonyls were visualized using the OxiSelect Protein Carbonyl Immunoblot Kit (Cell Biolabs). *Tnrf1*^-/-^ BMDMs were seeded in 24-well plates at 2.5x10^5^ one day prior to infection with indicated *L. pneumophila* strains at an MOI of 50 in the presence or absence of 10 µM H_2_O_2_. One hour after infection, cells were washed 3x with PBS^-/-^, media were replaced and cells were incubated for an additional 3 h in the presence or absence of H_2_O_2_. Cells were washed in ice-cold PBS and lysed in 120 µL ice-cold NP-40 lysis buffer (see above). Lysates were diluted in 3x Laemmli sample buffer and boiled for 10 min. Proteins were separated on a 4-20% gradient SDS-PAGE gel and transferred to PVDF membrane using a BioRad TransBlot semi-dry transfer apparatus. Membranes were processed according to manufacturer’s instructions using 10 min wash steps. Blots were stripped and re-probed with rabbit α-β-actin antibody (1:1000; Cell Signaling Technology) followed by goat-α-rabbit-HRP (1:5000; ThermoFisher). Blots were visualized as described above and densitometry was performed using Fiji ImageJ and Adobe Photoshop software.

### Yeast Two-Hybrid Analysis

Yeast two-hybrid screening was performed by Hybrigenics Services, S.A.S., Evry, France (http://www.hybrigenics-services.com). The *legC4* coding sequence was PCR-amplified from pSN85::*legC4* [20] and cloned into pB66 as a C-terminal fusion to Gal4 DNA-binding domain (Gal4-C4). The construct was checked by sequencing and used as a bait to screen a random-primed Mouse Spleen library constructed into pP6. pB66 derives from the original pAS2ΔΔ vector [102] and pP6 is based on the pGADGH plasmid [103].

Sixty million clones (6-fold the complexity of the library) were screened using a mating approach with YHGX13 (Y187 ade2-101::loxP-kanMX-loxP, mat-α) and CG1945 (mat-a) yeast strains as previously described [102]. 303 His+ colonies were selected on a medium lacking tryptophan, leucine and histidine, and supplemented with 2mM 3-aminotriazole to handle bait auto-activation. The prey fragments of the positive clones were amplified by PCR and sequenced at their 5’ and 3’ junctions. The resulting sequences were used to identify the corresponding interacting proteins in the GenBank database (NCBI) using a fully automated procedure. A confidence score (PBS, for Predicted Biological Score) was attributed to each interaction as previously described [104].

### Immunofluorescence Microscopy

To quantify LAMP1+ LCVs in naïve macrophages, 1x10^5^ BMDMs were seeded on poly-L-lysine (PLL)-coated glass coverslips in 24-well plates and infected with *L. pneumophila* at a multiplicity of infection (MOI) of 25 in triplicates. At 1h post-infection, cells were washed 3x in PBS to remove extracellular bacteria and cells were incubated for an additional 8 h in fresh media was added. Quantification of LAMP1+ LCVs in rTNF-treated BMDMs was performed as described but media were supplemented with 25 ng mL^-1^ rTNF throughout the course of infection, which was allowed to proceed for an additional 5 h after washing in PBS. Coverslips were fixed with 4% paraformaldehyde (PFA; Thermo Scientific), permeabilized in ice cold methanol, and blocked with blocking buffer [0.1% saponin (w/v), 1% HIFBS (v/v), 0.5% bovine serum albumin (BSA; w/v) in PBS]. Coverslips were stained with 1:1,000 rabbit α-*L. pneumophila* (Invitrogen, PA17227), 1:2,000 rat α-LAMP1 (Developmental Studies Hybridoma Bank) primary antibodies and 1:500 Alexa 488-conjugated goat α-rabbit and Alexa594-conjugated goat α-rat secondary antibodies (ThermoFisher) in blocking buffer. Nuclei were stained with Hoechst (ThermoFisher) at 1:2,000. Coverslips were mounted on glass slides with ProLong Gold Antifade Mountant (ThermoFisher). LAMP1+ LCVs were scored on a Leica DMiL LED inverted epifluorescence microscope (*n*=300 cells/strain) and the researcher scoring the samples were blind to their identity.

For confocal microscopy of transfected cells, HeLa cells were seeded on PLL-coated coverslips and transfected as described above. Coverslips were fixed in 4% PFA and processed as described above using mouse α-FLAG M2 (Sigma; 1:1,000) and rabbit α-GFP (Abcam; ab6556; 1:5,000) primary antibodies and Alexa546-conjugated goat α-mouse and Alexa488-conjugated goat α-rabbit antibodies (ThermoFisher; 1:500). Images were captured on a Zeiss LSM-5 PASCAL laser scanning confocal microscope (KSU Division of Biology Microscopy Facility). Co-localization was scored using the Pearsons algorithm for individual cells (*n*=10) using the colocalize tool Fiji ImageJ software.

For confocal microscopy of infected macrophages, 1.5x10^5^ cells were seeded on PLL-coated coverslips and infected as described above. Coverslips were fixed with 4% PFA and processed as above using rabbit α-*L. pneumophila* (Millipore Sigma; 1:1000) and mouse α-PA28α (SantaCruz; 1:100) primary antibodies, Alexa488-conjugated goat α-rabbit and Alexa594-conjugated goat α-mouse secondary antibodies, and Hoechst (ThemoFisher; 1,2000). Images were captured using an Olympus FluoView FV1000 confocal Laser scanning microscope (MSU Center for Advanced Microscopy).

### *L. pneumophila* intracellular growth curves

To quantify *L. pneumophila* intracellular replication, macrophages were seeded in 24-well tissue culture plates and infected in triplicates at an MOI of 1 the next day. Differentiated BMDMs were seeded at 2.5x10^5^ in seeding media (RPMI, 10% HIFBS, 7.5% L929 cell supernatant), THP-1 cells were seeded at 5x10^5^ and differentiated with 100 nM PMA in RPMI 10% HIFBS for three days. Cells were washed with PBS and incubated in the absence of PMA for one day prior to infection. Stable RAW 264.7 cells were seeded at 2.5x10^5^ in DMEM 2.5%HIFBS. At 1 h post-infection, cells were washed 3x with PBS followed by addition of fresh media. Macrophages were lysed in sterile water and colony forming units (CFU) were enumerated at the indicated time points as described [20]. Where indicated, 25 ng mL^-1^ rTNF (Gibco), 20-50 µM H_2_O_2_ (VWR), 10 nM bafilomycin A1 (BAF; ApexBio), or 1 µM MCC950 (ApexBio) were added at the time of infection and maintained throughout. DMSO was used as a vehicle control where indicated.

### Enzyme-linked immunosorbent assay (ELISA)

To quantify TNF secretion, 2.5 x 10^5^ BMDMs were seeded in 24-well tissue culture plates and infected with the indicated *L. pneumophila* strains at an MOI of 10. After 1 h of infection, cells were washed with PBS, and media were replaced. Supernatants were at 8 h post-infection and used fresh or stored at -20°C for up to 1 week and TNF was quantified using mouse TNF-α ELISA MAX kit (BioLegend) following manufacturer’s instructions. Absorbance at 450 nm was quantified on a BioTek Epoch2 microplate reader.

### Cytotoxicity assay

To quantify cell death, lactate dehydrogenase (LDH) activity in cell supernatants was quantified. BMDMs were seeded in triplicates at 2.5x10^5^ in a 24-well tissue culture plate for 24 h and infected with the indicated *L. pneumophila* strains at an MOI of 10 in 500 µL of seeding media (RPMP/10% HIFBS/7.5% L929 cell supernatant) and incubated for the indicated times. Supernatants were transferred to a 96-well plate and centrifuged at 200 r.c.f. for 10 min. LDH was quantified using the CytoTox 96 Non-Radioactive Cytotoxicity Assay (Promega) according to manufacturer’s instructions. Absorbance at 490 nm was quantified on a BioTek Epoch2 microplate reader and percent cytotoxicity was calculated by normalizing absorbance values to cells treated with lysis buffer. Where indicated, cells were primed with 1 µM PAM_3_CSK_4_ (Tocris) for 24 h prior to infection and plasmid expression of *legC4* was induced with 1 mM IPTG.

### Statistical analysis

Statistical analyses were performed with GraphPad Prism 9 software using either Students’ *t*-test, or two-way ANOVA, as indicated, with a 95% confidence interval. Data are presented as mean ± standard deviation (s.d.) or standard error of the mean (s.e.m.) and, unless otherwise indicated, statistical analyses were performed on samples in triplicates.

## Acknowledgements

We thank Dr. Russell Vance for femurs from *Casp1*^-/-^ mice and helpful discussions. We also thank Andrew Haskell for assistance within blinded microscopic scoring of *L. pneumophila*-infected cells. This work was funded by NIH/NIGMS P20GM130448 (to S.R.S.); Kansas-INBRE (K-INBRE) Postdoctoral Fellowship P20GM103418 (to D.C.) and Semester Scholar Award (to A.G.S.); Kansas State University Johnson Cancer Research Center Summer Stipend Award (to T.N.) and Faculty Expansion Award (to S.R.S.); and startup funds from Kansas State University and Michigan State University (to S.R.S.)

