## Supplemental Figures for "Effector-mediated subversion of proteasome activator (PA)28αβ enhances host defense against *Legionella pneumophila* under inflammatory and oxidative stress conditions"

**Supplementary Information (SI)**

### SI Figures

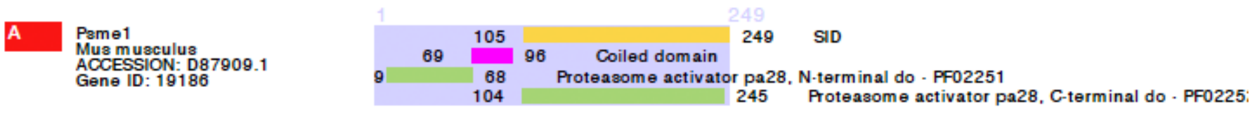


**Figure S1. All clones of PA28α shown to bind LegC4 by Y2H contained the C-terminal domain.** Schematic diagram of yeast two-hybrid results showing mouse PA28α (*Psme1*) domains and the selected interaction domain (SID) from amino acid residues 105-249.

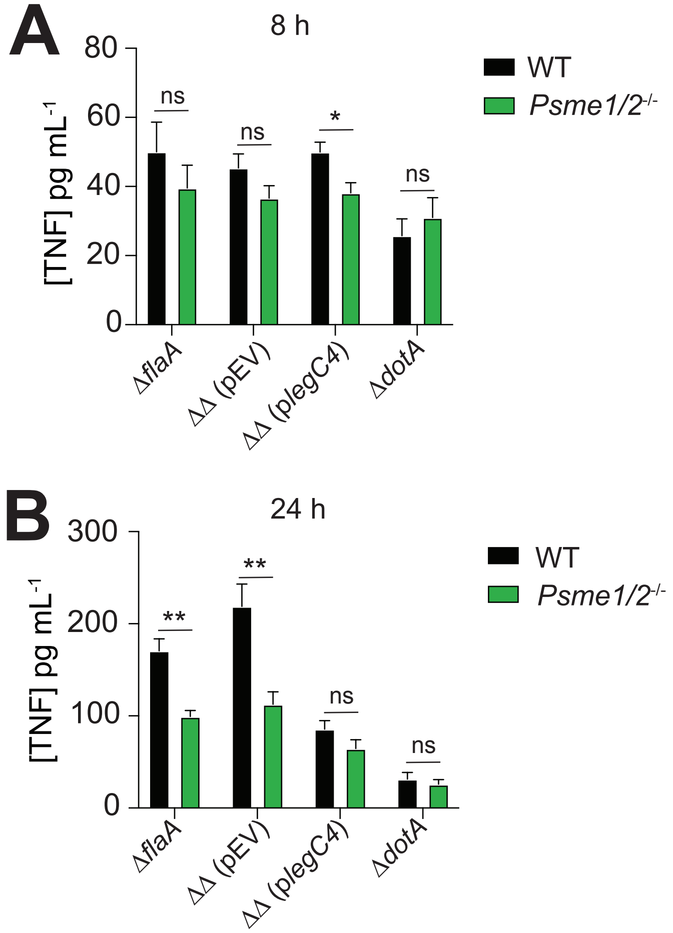


**Figure S2. TNF secretion from *L. pneumophila*-infected BMDMs is not increased by LegC4 or loss of PA28αβ.** TNF WT or *Psme1/2*$^{-/-}$ BMDMs infected for **(A)** 8 h or **(B)** 24 h with *L. pneumophila* Δ*flaA*, Δ*flaA*Δ*legC4* (pEV), Δ*flaA*Δ*legC4* (p*legC4*), or the avirulent Δ*dotA* control at a multiplicity of infection of 10. Plasmid expression of *legC4* was induced with 1 mM IPTG. Data shown are mean ± s.d. on samples in triplicates for a single experiment and are representative of results from three independent experiments. Asterisks denote statistical significance by two-way ANOVA (**P*<0.05; ***P*<0.05). ns; not significant.

.

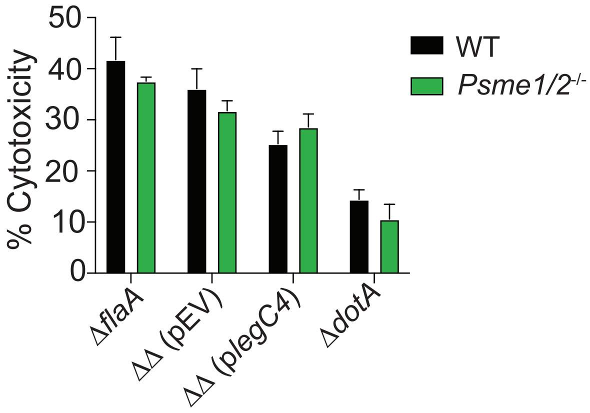


**Figure S3. Neither LegC4 nor loss of PA28αβ impair viability of *L. pneumophila*-infected BMDMs.** WT or *Psme1/2*-/- BMDMs were infected in triplicates with *L. pneumophila* strains (MOI of 10) and LDH in cell supernatants was quantified at 10h post-infection. Percent cytotoxicity was calculated by normalizing absorbance values to a lysis control (100% cytotoxicity). UI, uninfected cells. Plasmid expression of *legC4* was induced with 1 mM IPTG. Data shown are mean ± s.d. of triplicate samples for a single experiment and are representative of results from three independent experiments.

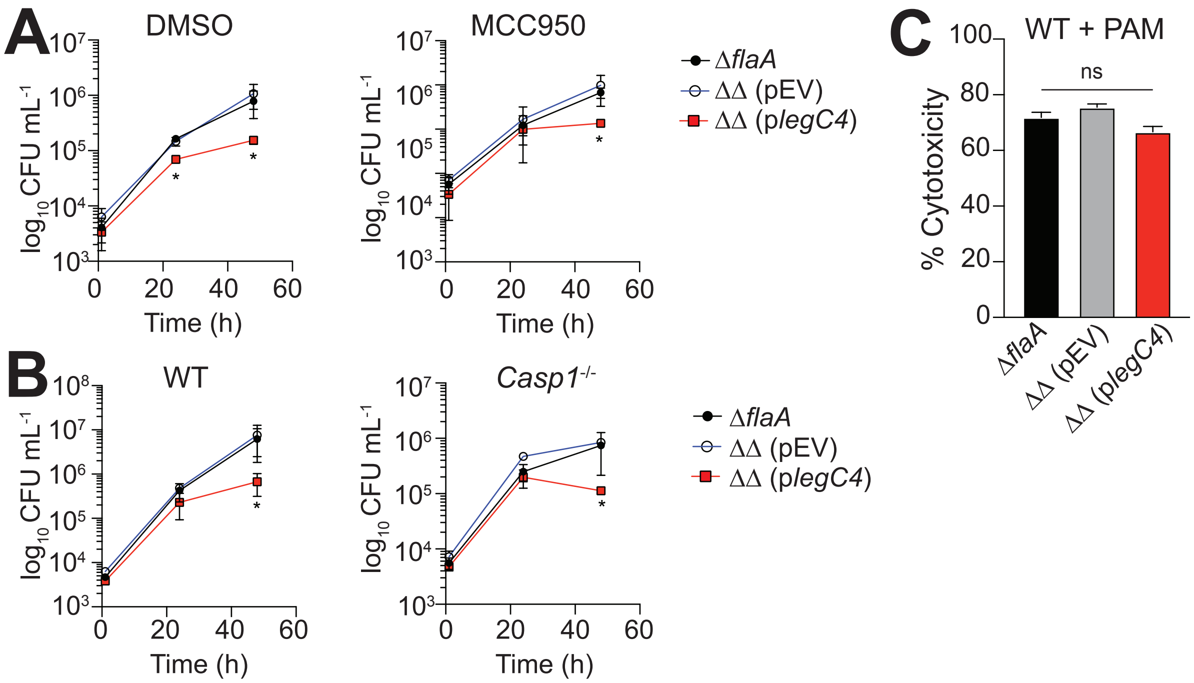


**Figure S4. LegC4-mediated restriction is not dependent on caspase-1 or NLRP3 inflammasome activity**. **(A)** WT BMDMs were infected with *L. pneumophila* strains (MOI of 1) in the presence of either 1μM MCC950 or volume equivalent of DMSO (vehicle) and CFU were enumerated at the indicated time points. **(B)** WT and *Casp1*$^{-/-}$ BMDMs were infected with *L. pneumophila* (MOI of 1) and CFU were enumerated at the indicated time points. Data are shown as mean ± s.d. of pooled results of two independent experiments with triplicate wells per condition in each experiment. Asterisks denote statistical significance by Two-way ANOVA (**P*<0.05; ***P*<0.01). Plasmid expression of *legC4* was induced with 1 mM IPTG. **(C)** WT BMDMs were primed with 1 μM $PAM_3CSK_4$ (PAM) for 24 h and infected with the indicated *L. pneumophila* strains at an MOI of 10 for 6 h and cytotoxicity was quantified by LDH release assay. Data shown are mean ± s.d. of triplicate samples for a single experiment and are representative of results from three independent experiments. Ns; not significant by Two-way ANOVA.

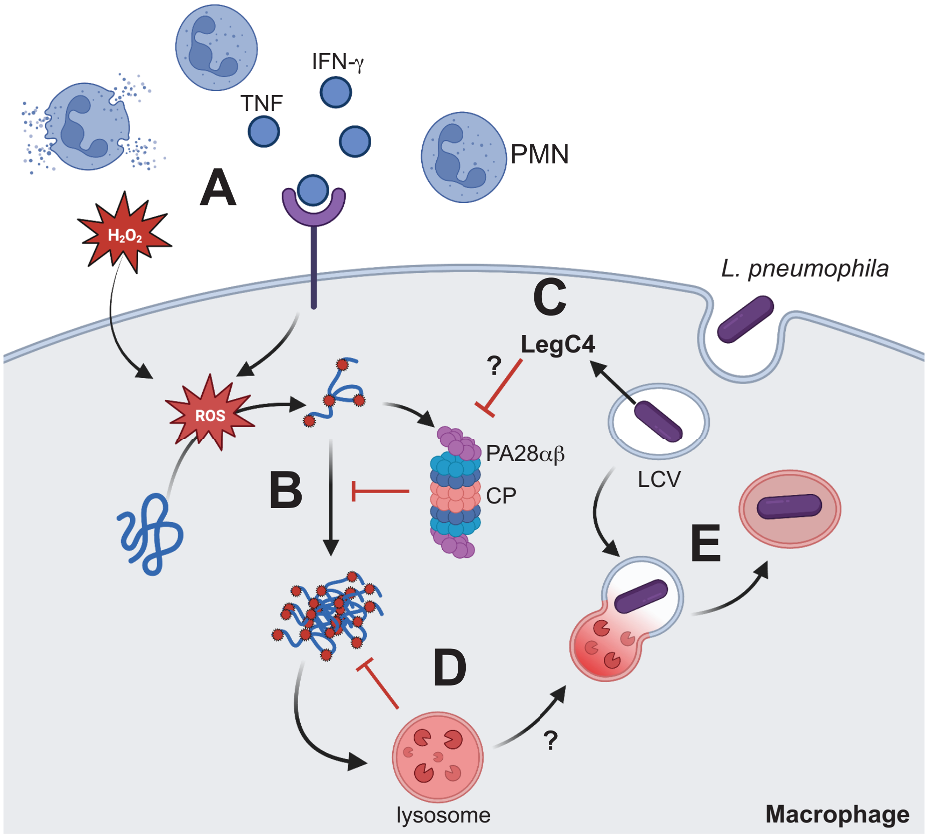


**Figure S5.** Schematic model of the LegC4 restriction mechanism. **(A)** TNF, IFN-γ and reactive oxygen species (ROS) are produced by infected and bystander immune cells induce oxidative stress in activated macrophages via intracellular ROS production and by passive diffusion of membrane-permeable extracellular ROS (H2O2). **(B)** ROS indiscriminately carbonylate amino acid side chains, which perturbs protein folding and function, and turnover of these damaged proteins is mediated by PA28ab-CP proteasomes. **(C)** LegC4 translocated into *L. pneumophila*-infected macrophages binds PA28a and modulates its activity by an unknown mechanism. **(D)** Impaired proteasome activity leads to formation of carbonylated protein aggregates, which are impervious to proteasomal degradation and trigger upregulation of lysosome biogenesis and fusogenic activity. **(E)** Increased phagolysosomal fusion with the *Legionella*-containing vacuole (LCV) may result from global increases in lysosomal degradation to maintain proteostasis under oxidative stress conditions. Image created with Biorender.com.

**SI Tables**

**Table S1**. Clones of proteasome regulators identified as LegC4 interacting partners by yeast two-hybrid.

See file: Table S1.xlsx

**Table S2.** *Legionella* strains used in this study

| Strain | Description | Resistance | Ref(s) |
|---|---|---|---|
| *Legionella pneumophila* SRS43 | | | |
| Δ*flaA* | FlaA-deficient parental strain | $Sm^R$ | (1) |
| Δ*flaA*Δ*legC4* | LegC4-deficient Δ*flaA* strain | $Sm^R$ | (1) |
| Δ*flaA*Δ*legC4* (pEV) | Harboring empty pSN85 vector | $Sm^R$, $Cm^R$ | (1) |
| Δ*flaA*Δ*legC4* (p*legC4*)[a] | Harboring pSN85::*legC4* | $Sm^R$, $Cm^R$ | (1) |
| Δ*flaA*Δ*legC4* (pJB) | Harboring empty pJB1806 vector | $Sm^R$, $Cm^R$ | (1) |
| Δ*flaA*Δ*legC4* (pJB*legC4*)[b] | Harboring pJB1806::p*legC4* | $Sm^R$, $Cm^R$ | (1) |
| Δ*dotA* | Avirulent control | $Sm^R$, $Cm^R$ | (2) |

a – gene expression induced with 1 mM IPTG
b – *legC4* expression from endogenous promoter

**Table S3.** Plasmids used in this study

| Plasmid | Description | Resistance | Ref(s) |
|---|---|---|---|
| *L. pneumophila* expression | | | |
| pSN85 (pEV) | *L. pneumophila* expression vector | $Cm^R$ | (3) |
| pSN85::*legC4* (p*legC4*)[a] | For expression of *3xflag-legC4* | $Cm^R$ | (1) |
| pJB1806 (pJB) | *L. pneumophila* expression vector | $Cm^R$ | (4) |
| pJB1806::*legC4* (pJB*legC4*)[b] | *legC4* expression from endogenous promoter | $Cm^R$ | (1) |
| *E. coli* expression | | | |
| pGEX6P1 | For expression of GST-fusion proteins | $Amp^R$ | GE Healthcare |
| pGEX::*psme1* | Expression of GST-PA28a | $Amp^R$ | This study |
| pGEX::*legC4* | Expression of GST-LegC4 | $Amp^R$ | This study |
| pGEX::*lgt1* | Expression of GST-Lgt1 | $Amp^R$ | (2) |
| pT7HMT | Expression of $His_6$-Myc fusion proteins | $Kan^R$ | (5) |
| pT7HMT::*psme1* | Expression of $His_6$-Myc-PA28a | $Kan^R$ | This study |
| pT7HMT::*legC4* | Expression of $His_6$-Myc-LegC4 | $Kan^R$ | This study |
| pT7HMT::*lgt1* | Expression of $His_6$-Myc-Lgt1 | $Kan^R$ | (2) |
| Mammalian Expression | | | |
| pcDNA 3FLAG 4/TO | For ectopic FLAG-fusion production | $Amp^R$ | (6) |
| pcDNA::*3xflag-legC4* | Ectopic production of FLAG-LegC4 | $Amp^R$ | This study |
| pEGFPC1 | Expression of GFP-fusion proteins | $Kan^R$ | Clontech |
| pEGFPC1::*psme1* | Ectopic production of GFP-PA28$\alpha$ | $Kan^R$ | This study |
| pCMV-3Tag-4a::*psme1*[c] | Ectopic production of PA28$\alpha$-Myc | $Kan^R$ | This study |

a – gene expression induced with 1 mM IPTG
b – *legC4* expression from endogenous promoter
c – purchased from Genscript (Piscataway, New Jersey)

**Table S4**. Oligonucleotide primers used in this study

| Name | Sequence (5'→3')[a] |
|---|---|
| LegC4BamHI-F3 | ATTGGATCCTTGATTCATTATGTATCCTTG |
| LegC4NotI-R | ATTGCGGCCGCTTATAGCTTAATATCAAAAG |
| Psme1Sal1-F | ATTGTCGACATGGCCACACTGAGGGTCCATCCC |
| Psme1BamHI-R | ATTGGATCCTCAATAGATCATTCCCTTGGTTTC |
| Psme1BamHI-F | ATTGGATCCATGGCCACACTGAGGGTCCATCCC |
| Psme1NotI-R | ATTGCGGCCGCATTCAATAGATCATTCCCTTGGTTTC |

a – restriction endonuclease cleavage sites are underlined.
